# Cold-inducible promoter-driven knockdown of *Brachypodium* antifreeze proteins confers freeze sensitivity

**DOI:** 10.1101/2022.02.15.480542

**Authors:** Collin L. Juurakko, Melissa Bredow, George C. diCenzo, Virginia K. Walker

**Affiliations:** Department of Biology, Queen’s University, Kingston, Ontario, Canada; School of Environmental Studies, Queen’s University, Kingston, Ontario, Canada

**Keywords:** *Brachypodium distachyon*, cold acclimation, freeze tolerance, abiotic stress, antifreeze proteins, ice-recrystallization inhibition

## Abstract

The model forage crop, *Brachypodium distachyon*, has a family of ice recrystallization inhibition (*BdIRI*) genes, which encode antifreeze proteins that function by adsorbing to ice crystals and inhibiting their growth. The genes were previously targeted for knockdown using a constitutive CaMV 35S promoter and the resulting transgenic *Brachypodium* showed reduced antifreeze activity and a greater susceptibility to freezing. However, the transgenic plants also showed developmental defects with shortened stem lengths and were almost completely sterile, raising the possibility that their reduced freeze tolerance could be attributed to developmental deficits. A cold-induced promoter from rice (pr*OsMYB1R35*) has now been substituted for the constitutive promoter to generate temporal miRNA-mediated *Brachypodium* antifreeze protein knockdowns. Although transgenic lines showed no apparent pleiotropic developmental defects, they demonstrated reduced antifreeze activity as assessed by assays for ice-recrystallization inhibition, thermal hysteresis, electrolyte leakage, leaf infrared thermography, and leaf damage after infection with an ice nucleating phytopathogen. Strikingly, the number of cold-acclimated transgenic plants that survived freezing at -8 °C was reduced by half or killed entirely, depending on the line, compared to cold-acclimated wild type plants. Although these proteins have been studied for almost 60 years, this is the first unequivocal demonstration in any organism of the utility of antifreeze protein function and their contribution to freeze protection, independent of obvious developmental defects. These proteins are thus of potential interest in a wide range of biotechnological applications from accessible cryopreservation, to frozen product additives, to the engineering of transgenic crops with enhanced freezing tolerance.

## 1 Introduction

Increased freezing events and altered freeze-thaw cycles in response to climate change present major challenges to agriculture with single frosts costing billions of dollars (Witney and Arpaia, 1991; Sinha and Cherkauer, 2010; Smith and Katz, 2013; Kreyling, 2019; NOAA, 2021; Smith et al., 2021). In the field, the formation of ice at high sub-zero temperatures is initiated by ice nucleation active (INA+) bacteria and is a major driver of crop destruction (Snyder and Melo-Abreu, 2005). Since they cannot escape low temperatures, many temperate climate plants have adopted a freeze tolerant strategy with some producing antifreeze proteins (AFPs) to help prevent freeze damage (Bredow and Walker, 2017; Juurakko et al., 2021a). In contrast, other organisms such as polar fish and temperate arthropods, which can escape low temperatures and find hibernacula, frequently adopt a freeze-avoidance strategy that can also employ AFPs, in this case to lower the freezing point relative to the melting point, also known as the thermal hysteresis (TH) gap (Duman, 2001; Bar Dolev et al., 2016; Kim et al., 2017). Despite the discovery of these proteins, first recognized almost 5 decades ago in *Tenebrio* beetles (Ramsay, 1964), until recently, there has been no formal evidence of their contribution to low temperature survival in any organism. This changed with the engineering of transgenic grass lines with knocked down expression of AFPs, resulting in greater freeze susceptibility (Bredow et al., 2016). However, important as these results were, it was worrying that the knockdowns were associated with other phenotypes including stunted growth and almost complete sterility, suggesting to naysayers that the greater freeze susceptibility could be due to unhealthy plants rather than the low AFP activity. To test that possibility, and to verify that AFPs do indeed make a crucial contribution to freeze survival, it was important that new transgenic lines be made that were unencumbered with detrimental phenotypes.

AFPs are also known as ice-binding proteins or ice-recrystallization inhibition (IRI) proteins. The plant freeze-tolerant overwintering strategy may be associated with AFPs that are characterised by low TH activity but a high IRI activity, which keeps ice crystals small even when the temperature fluctuates near 0 °C. This is important since ice forms in the apoplast, frequently due to INA+ bacteria, such as *Pseudomonas syringae*, gaining entry through stomatal openings, hydathodes, or wounding sites which themselves may result from surface tissue ice formation (Lindow et al., 1982; Ashworth and Kieft, 1995; Wisniewski and Fuller, 1999; Pearce and Fuller, 2001). Some plant AFPs can even attenuate the INA+ activity of *P. syringae*, possibly by binding to the bacterial ice nucleating proteins (INPs) and resulting in a modest lowering of the freeze temperature (Tomalty and Walker, 2014; Bredow et al., 2018). The apoplast has a lower solute concentration and thus freezes before other tissues. Thus, in order to combat catastrophic freezing, AFPs are produced and secreted to the apoplast (Griffith et al., 1992; Marentes et al., 1993; Hon et al., 1994; Hon et al., 1995; Antikainen et al., 1997; Bredow et al., 2016). The property of AFPs to irreversibly adsorb to ice crystals is key to membrane protection and explains why these proteins can be employed in both freeze-tolerant and freeze-avoidant strategies. In the absence of other ice management mechanisms, uncontrolled ice growth in the apoplast can lead to cellular death by dehydration through exclusion of solutes or the piercing of membranes, thus presenting the primary battleground between AFPs and ice (Lindow et al., 1982; Melo-Abreu et al., 2016).

*Brachypodium distachyon* (hereinafter *Brachypodium*) contains 7 genes encoding AFPs (*BdIRI1-7*). Their translation products are hydrolysed, likely in the apoplast, to generate two independent proteins, a leucine-rich repeat (LRR) protein that has not been characterised, and an AFP (Bredow et al., 2016). The *BdIRI* gene sequences are sufficiently similar so that a single miRNA could be designed to attenuate the translation of all 7 corresponding mRNAs with no obvious off-target binding (Bredow et al., 2016). As noted, previously generated transgenic *Brachypodium* lines bearing the miRNA sequence, driven by the constitutive CaMV 35S promoter, were more susceptible to freeze damage than non-transgenic controls. Thus, although these experiments clearly connected AFPs with freeze protection, the lines also showed developmental deficits. We hypothesised that these secondary phenotypes were due to the constitutive promoter selected for those experiments, and that a cold-induced promoter, perhaps more similar to the native *BdIRI* promoters, would circumvent this problem and allow a fuller characterisation of newly-generated AFP knockdown lines. This has now been achieved and here we report an exploration of *BdIRI* regulation as well as showing that the experimental temporal attenuation of AFP expression is inextricably linked to greater freeze susceptibility.

## 2 Methods

### 2.1 Bioinformatic Analysis

*BdIRI* gene and protein sequences were retrieved from NCBI using up-to-date accessions (December 2020) using a BLAST search with the published proteins as queries (Bredow et al., 2016). The lack of data for the identification of suitable low-temperature inducible promoters in *Brachypodium* prompted the selection of the 1961 bp promoter associated with the rice, *Oryza sativa*, gene *OsMYB1R35*, which is induced in its host plant after cold stress (Li et al., 2017). The sequence was retrieved from the publicly available *O. sativa* genome on NCBI (accessed October 2017) based on the primer sequences described elsewhere (Li et al., 2017). The 1961 bp fragment was synthesized by GeneART (Thermo Fisher Scientific, Waltham, MA) with appropriate flanking restriction enzyme recognition sites. Conceptually translated *BdIRI* sequences were aligned and phylogenies prepared using the Clustal Omega Multiple Sequence Alignment tool (https://www.ebi.ac.uk/Tools/msa/clustalo/) (Sievers et al., 2011). Inter-domain hydrolytic cleavage sites in the amino acid sequence corresponding to the *BdIRI*s were predicted using the ExPASy PeptideCutter tool (https://web.expasy.org/peptide_cutter/; Gasteiger et al. 2005). Chromosomal positioning of the *BdIRI* genes was determined using NCBI’s genome browser. InterProScan (Version 83.0; https://www.ebi.ac.uk/interpro/) was used for the *in silico* prediction of the LRR and AFP domains in the *BdIRI* translated sequences (Blum et al., 2021). The Phyre2 Protein Fold Recognition Server (http://www.sbg.bio.ic.ac.uk/~phyre2/) was used for sequence-based homologous protein structure prediction of the corresponding *BdIRI* domains (Kelley et al., 2015). The SignalP 5.0 Server (http://www.cbs.dtu.dk/services/SignalP/) was used to forecast putative amino-terminal secretory signal peptide sequences (Armenteros et al., 2019). The DeepLoc 1.0 eukaryotic protein subcellular localization predictor (http://www.cbs.dtu.dk/services/DeepLoc-1.0/) was used to confirm the subcellular localization of proteins secreted to the apoplast (Armenteros et al., 2017).

### 2.2 Prediction of miRNA Targets and Analysis of Regulatory Elements

Identification of post-transcriptional regulation via endogenous miRNAs was performed on the *BdIRI* mRNA sequences using the psRNATarget miRNA prediction tool (http://plantgrn.noble.org/psRNATarget/) (Dai et al., 2018). The tool was used with data from miRBase (Release 21, June 2014) (Griffiths-Jones, 2004; Griffiths-Jones et al., 2006; Griffiths-Jones et al., 2007; Kozomara and Griffiths-Jones, 2010; Kozomara and Griffiths-Jones, 2014; Kozomara et al., 2019).

Upstream sequences to the translational start codons of selected genes were retrieved from NCBI (*B. distachyon* genome assembly v3 2020) in addition to the promoter sequence of p*rOsMYB1R35* from *O. sativa* rice (Li et al., 2017). Sequences were submitted to PlantCARE (http://bioinformatics.psb.ugent.be/webtools/plantcare/html/) (Lescot et al., 2002) for *cis*-regulatory element prediction and analysis. Raw output files were converted to text and input into excel where they were compiled for a comparative analysis of *cis*-regulatory elements among the putative promoter sequences used. *Cis*-regulatory elements specifically related to cold signalling and cold stress were manually annotated for putative promoter sequences of all *BdIRI*s and *OsMYB1R35*, designated as pr*BdIRIs1-7* and pr*OsMYB1R35*, respectively.

### 2.3 Plasmids Construction and *Brachypodium* Transformation

The GeneART pUC plasmid containing the pr*OsMYB1R35* promoter sequence was transformed into *Escherichia coli* DH5α cells (Thermo Fisher Scientific). The promoter was liberated from purified plasmid using *Bam*HI and *Bgl*II restriction enzyme digests for the 5’ and 3’ end, respectively. It was then ligated into the plant expression plasmid pCambia1380 (Marker Gene Technologies Inc., Eugene, OR, USA) and transformed into *E. coli* DH5α. A plasmid bearing a sequence corresponding to the miRNA (Bredow *et al*., 2016) with a 5’ *Bgl*II site, was amplified by polymerase chain reaction (PCR) and a 3’ *Spe*I restriction site was added with the use of appropriate primers. Similarly, the sequence encoding enhanced green fluorescent protein (eGFP; GenBank Accession no. U57607) was PCR-amplified and 5’ *Bgl*II and 3’ *Spe*I restriction sites were added. The pCambia1380 with the pr*OsMYB1R35* insert was then digested using *Bgl*II and *Spe*I enzymes and the purified amplified products, miRNA and eGFP, were ligated separately to create the plasmids pCambia1380:pr*OsMYB1R35*:miRNA and pCambia1380:pr*OsMYB1R35*:eGFP, respectively. These plasmids were independently transformed into *E. coli* DH5α and the veracity of their sequence subsequently confirmed by Sanger sequencing (CHU de Québec-Université Laval, Quebec City, QC, CA). DNA from the verified plasmids was isolated, purified and transformed into *Agrobacterium tumefaciens* (AGL1, Invitrogen, Carlsbad, CA, USA) (hereinafter, *Agrobacterium*). Again, the authenticity of the target sequences was checked by Sanger sequencing. AGL1 cells containing the confirmed plasmids were then used for transformation into *Brachypodium*.

Transgenic *Brachypodium* lines were generated using a modified method from Fursova *et al*. (2012) as follows. Transformed *Agrobacterium* cultures (50 mL), were grown in Luria Bertani (LB) broth containing 50 mg L^-1^ kanamycin to an OD_600_ of 1, pelleted at 5000 × *g* for 10 min, and washed in equal volumes of 2-(N-morpholino)ethanesulfonic acid (MES) infiltration buffer (10 mM MES, 10 mM MgCl_2_, pH 5.6). Pellets were resuspended in 50 mL of infiltration buffer containing 50 μM acetosyringone, 0.01% Silwet-L77 organosilicone surfactant, and leaf extracts containing phenolic metabolites to initiate efficient Ti plasmid transfer made from Australian tobacco, *Nicotiana benthamiana* leaves, rather than *Nicotiana tabacum* (Fursova *et al*., 2012). To prepare the tobacco extracts, up to 50 g of leaf tissue was harvested from six-week-old plants, cut into 1-3 cm^2^ squares, and incubated in 300 mL of infiltration buffer for 2 h. The liquid was removed by pressing a sterile beaker of slightly smaller diameter into the slurry and subsequently filter-sterilising the recovered liquid through 0.22-micron syringe filters (Thermo Fisher Scientific). Acetosyringone and surfactant were added following filter sterilisation and used to resuspend the washed *Agrobacterium* pellet. In parallel, mature wild type seeds (50 per trial) were harvested, surface-sterilised as described, and aseptically trimmed using a sterile scalpel to remove the upper quarter of the seed where the awn had been attached. The lower portion of the sterilised and trimmed seeds, with exposed embryos, were immediately added to the now-primed *Agrobacterium* culture and co-cultivated for 30 h at 21 °C with shaking at 200 rpm, in the dark.

Following co-cultivation, seeds were washed in infiltration buffer alone and plated on Linsmaier and Skoog (LS) agar (Phytotech Labs, Lenexa, KS, USA) containing 225 mg L^-1^ timentin, a wide-spectrum antibacterial, in order to kill residual *Agrobacterium*. The plates were sealed with micropore tape and put at *Brachypodium* standard growth conditions (see below). One week following successful germination, surviving T_0_ generation seedlings were sown to soil and grown. Hygromycin selection media was not used, since initial experiments suggested that this treatment was often detrimental to the seedlings. DNA was subsequently extracted from T_0_ plants using Monarch DNA extraction kits (New England Biolabs, Ipswich, MA, USA) and sequence confirmed (CHU de Québec-Université Laval, Quebec City, QC, Canada). Once T_0_ plants were brought to senescence, seeds were counted and harvested. Seeds from these plants (10 per line) were sterilised and germinated on LS agar with hygromycin. Selected, germinated seeds were sown in soil and brought to senescence representing the T_1_ generation. After collection, seeds (24 per line) were sterilised and germinated on LS agar without antibiotics. These T_2_ generation plants were used for rapid genotyping (see below) to determine homozygosity for the T_3_ generation without the need for selection.

### 2.4 Genotyping *Brachypodium* Transgenic Lines

Genotyping was done using a modified method from Ben-Amar et al. (2017). Briefly, seeds (24 per line) were sterilised, germinated, and sown in soil. Leaf tips (∼ 2 mm long) were cut at two weeks and placed in individual wells of 96 well PCR plates, each holding 50 µL of TE buffer at pH 8.0. The leaf tissue was gently ground using sterile fine-tip forceps, which were washed in 70% ethanol between samples. Once all samples were homogenised, plates were incubated at 60 °C for 10 min and then gently vortexed. Extracted samples (2 µL) from each line were combined into a 1.5 mL tube and gently vortexed, and from this pool containing extracts from each plant, 1 µL was used as the DNA template for a PCR screen of the insert. Lines contributing to the pooled extracts showing an amplified DNA band of an appropriate size after gel electrophoresis were then selected for individual sample screening. For this, 1 µL of sample extracts from individual plants were used as PCR templates, with lines showing a positive insert band for every individual considered to be homozygous. The reference gene, S-adenosylmethionine decarboxylase (*SAMDC*) was used as a PCR control.

### 2.5 Western Blot Analysis

Western blot analysis was conducted on leaf samples from three-week-old wild type (Bd21) plants or transgenic plants maintained under standard conditions (non-acclimated; NA) or transferred to 4 °C for 48 h (cold-acclimated; CA, see details below). These plants were obtained by using seeds from wild type or transgenic pr*OsMYB1R35*:eGFP plants. Subsequent to surface sterilisation, the seeds were germinated on selection media with 50 mg mL^-1^ hygromycin B (BioShop Canada Inc., Burlington, ON, Canada). After sowing on potting soil, they were grown for three weeks before harvest of 200 mg of leaf tissue from each plant. After freezing with liquid nitrogen and grinding to a fine powder using a sterile micro pestle, the samples were resuspended in extraction buffer (5 mM DTT, 1% Sigma P9599 protease inhibitor, 0.1% Igepal, 2 mM NaF, 1.5 mM activated Na_3_VO_4_, 0.5 M Tris-HCl pH 7.5, 10% glycerol, 0.15 M NaCl) and shaken at 150 rpm at 4 °C for 4 h. Following centrifugation (13,000 x *g* at 4 °C for 30 min), the protein concentration in the supernatants was estimated (Bradford reagent, Thermo Fisher Scientific) and then diluted so that samples were equivalent. Subsequently, 100 µL of the samples were denatured by the addition of 50 µL of Laemmli Sample Buffer (45% glycerol, 10% SDS, 0.5 M Tris pH 6.8, 0.045% w/v bromophenol blue, 0.006% 1 M DTT) and boiled for 5 min. Samples were run using a semi-dry transfer apparatus (Bio-Rad Laboratories, Hercules, CA, USA) following the manufacturer’s recommended protocols. The membrane was blocked for 1 h using a 5% (w/v) skim milk powder in Tris-buffered saline with 0.1% Tween^®^ 20 detergent (TBST) while shaking at room temperature.

GFP was detected on the membranes using anti-GFP mouse-IgG monoclonal (clones 7.1 and 13.1) antibody (Roche, Basel, Switzerland) in a 1/500 dilution in 5% (w/v) skim milk powder TBST solution with gentle shaking overnight at 4 °C in the dark. The secondary antibody, anti-mouse IgG peroxidase-conjugated (Sigma-Aldrich, St. Louis, MO, USA) was used in a 1/4000 dilution in the same skim milk-TBST solution with gentle shaking at room temperature for 1 h. Coomassie brilliant blue stained ribulose-1,5-bisphosphate carboxylase-oxygenase (better known as RuBisCO) large chain (RbcL) was used as a 55 kDa loading control. Blots were washed with TBST buffer three times for 10 min each time and inserted between acetate transparency sheets and imaged on a ChemiDoc Touch Imaging System (Bio-Rad Laboratories) using Immobilon western chemiluminescent HRP substrate (MilliporeSigma, Boston, MA, USA). Western blot analysis was done using Image Lab Software (Bio-Rad Laboratories). Purified recombinant eGFP was used as a positive control and western blots were repeated in triplicate.

### 2.6 Plant Material and Growth Conditions

*Brachypodium* seeds of an inbred line, ecotype Bd21 (RIKEN, Tokyo, Japan), were prepared by soaking in sterile water for 1 h followed by careful removal of the lamella, awn, and any remaining appendages still attached to the harvested floret. Seeds were washed in a 40% bleach, 0.04% w/v Silwet-L77 solution for 4 min followed by a 2 min wash in 70% ethanol and 4 washes in sterile water and finally dried with filter paper soaked in 100% ethanol. Seeds were transferred to LS agar plates using sterilised forceps and the plates were subsequently sealed using sterile micropore tape. All seed work was done in a UV sterilised laminar flow hood. Plates were subsequently wrapped in aluminium foil and placed at 4 °C to synchronise germination for four days.

After this time, plates were moved to a climate-controlled growth chamber (Conviron CMP4030, Controlled Environments Limited, Winnipeg, MB, Canada) at standard *Brachypodium* growth conditions of 70% relative humidity and 24 h cycles of 20 h light (∼150 μmol m^−2^s^−1^) at 24 °C followed by 4 h with no light at 18 °C. After one week, seeds were transplanted to pots filled with moist potting soil and then fertilised bi-weekly using 10-30-20 Plant-Prod MJ Bloom (Master Plant-Prod, Brampton, ON, Canada). Prior to assaying, CA plants were moved to a separate chamber (Econair GC-20, Ecological Chambers Inc., Winnipeg, MB, Canada) maintained at 4 °C where they were subjected to a shortened day cycle of 6 h of light (∼150 μmol m^−2^s^−1^) and 20 h of darkness for 48 h. NA plants remained at standard conditions.

### 2.7 Crude Lysate and Apoplast Extract Preparations

AFP activity was assayed in extracts prepared as described previously (Bredow et al., 2016). After CA, 50 mg of leaf tissue was taken from the three-week-old plants and ground to a fine powder using a sterile pestle after being flash frozen with liquid nitrogen. After suspension in 400 µL of NPE buffer (25 mM Tris, 10 mM NaCl, pH 7.5, EDTA-free protease inhibitor tablets), the slurry was shaken for 4 h at 4 °C in the dark on a GyroMini nutating mixer (Labnet International, Edison, NJ, USA). Samples were centrifuged at 13,000 × *g* for 5 min and placed at 4 °C for 5 min and then the centrifugation and the incubation were repeated. The supernatant was transferred to 1.5 mL tubes and centrifuged again (13,000 × *g* for 5 min) before returning to 4 °C. Protein concentration was estimated using a Nanodrop One (Thermo Fisher Scientific; using a mass extinction coefficient (ε_1%_) of 10 at 280 nm for 10 mg mL^-1^ with a baseline correction at 370 nm, as recommended by the manufacturer). Readings were performed in triplicate for each sample, with samples routinely diluted to 1 mg mL^-1^ unless stated otherwise.

Apoplast extracts were prepared as previously described (Pogorelko et al., 2011) with minor modifications. Briefly, 0.5 g of leaf tissue of three-week-old NA and CA wild type and knockdown lines were collected. Leaf tissue was sliced with a sterile scalpel into 1 cm segments and placed vertically into a 10 mL syringe with the tip sealed with parafilm. Chilled, 4 °C extraction buffer (5 mL of 25 mM Tris–HCl, 50 mM EDTA, 150 mM MgCl_2_, pH 7.4) was added and the syringe was placed under vacuum for 1 min, four separate times, until bubbles ceased and leaves were fully infiltrated and darker in appearance. Excess buffer was drained, and leaf tissue was transferred to a 3 mL syringe. The syringe, without the plunger, was then placed into a 15 mL conical tube and centrifuged at 1000 x *g* for 10 min at 4 °C. Apoplast fluid was collected at the bottom of the conical tube and transferred to a fresh 1.5 mL microcentrifuge tube. Protein concentration was estimated using a Synergy H1 microplate reader (BioTek Instruments, Inc., Winooski, VT, USA) with a Take3 Micro-Volume Plate (BioTek Instruments) at A_280_, as described for the lysates. Samples were normalised and diluted as described prior to assaying. All work was carried out at 4 °C.

### 2.8 AFP assays

To assess IRI activity, a modified “splat” assay was performed as previously described (Bredow et al., 2017). Briefly, samples (10 µL) were pipetted 1 m above a glass cover slip, equilibrated on an aluminium block that had been chilled with dry ice for 1 h, to ensure the formation of a thin layer of small ice crystals prior to transfer into a hexane-containing bath for annealing maintained at -6 °C. Images were captured through cross-polarising films at 10x magnification, immediately after transfer to the bath and again after annealing for 18 h. Lysates and apoplast extracts were subjected to a standardised dilution series and assays were done a minimum of three times for all samples.

Samples for TH assays were prepared as described (Bredow et al., 2020), but with 200 mg of leaf tissue and 800 µL of buffer. Amicon Ultra-0.5 micro-concentrators (Millipore) were used to concentrate supernatants 4-fold after centrifugation (13,000 x *g* for 10 min). TH was determined on a nanoliter osmometer as described (Middleton et al., 2014). Ice crystal morphology was recorded during the TH assays with images captured using a microscope video camera at 50x magnification and in triplicate.

### 2.9 Electrolyte Leakage, Infrared Thermography, Whole Plant Freezing Assays and Infections

Electrolyte leakage assays were conducted as described (Bredow *et al*., 2016). Briefly, leaf tips (∼ 4 cm long) were excised from three-week-old plants and placed individually into sterilised glass test tubes containing 100 µL of deionized water with the wounded end up and the tip submerged in the water. Sterilised foam plugs were used to ensure that the leaf tips remained submerged. One set of control tubes contained leaf tips from individual, numbered plants and were kept at 4 °C in the dark. The second companion set of tubes with leaves were placed in a programmable circulating ethylene glycol temperature-controlled bath set at 0 °C. The temperature was ramped down from 0 °C to -1 °C over 30 min and then a single ice chip was added to initiate ice crystal growth, with the temperature then lowered 1 °C every 15 min until the final freezing temperature of -10 °C was reached. The tubes were then placed at 4 °C in the dark with the control tubes and left overnight. After transfer of the tube contents to 50 mL centrifuge tubes containing 25 mL of deionized water, they were shaken horizontally on a G2 Gyrotory Shaker (New Brunswick Scientific, Edison, NJ, USA) at 150 rpm for 18 h in the dark at room temperature. Initial conductivity (C_i_) was measured using an Oakton CON 510 conductivity metre (OAKTON Instruments, Vernon Hills, IL, USA) prior to autoclaving the samples for 45 min. Samples were cooled to room temperature overnight and final conductivity (C_f_) was then measured. The percentage of electrolyte leakage was calculated using (100 × C_i_ / C_f_). The assay was performed in triplicate using leaves from 10 individual plants per line and condition.

To visualise ice propagation in leaves and the influence of AFPs, infrared thermography was used, which detects emitted infrared energy provided that the objects have sufficiently high levels of emissivity over background. In plants, emissivity ratings are typically in the range of 0.98 (López et al., 2012; Chen, 2015). To enhance the contrast, household aluminium foil was used as a background because of its low emissivity of 0.05 (Qin et al., 2017). The FLiR One Pro – iOS (FLIR Systems, Wilsonville, OR, USA) with Vernier Thermal Analysis Plus application (Apple App Store) (Vernier Software and Technology, Beaverton, OR, USA) was used to capture thermography data. *Brachypodium* leaves of equal length (∼2.5 cm) were freshly excised from CA three-week-old plants and placed on a stage lined with aluminium foil touching the surface of a circulating ethylene glycol bath set at 1 °C. Leaf tissue was annealed for 30 min before distilled water (10 µL) was pipetted onto the wounded end of the leaf and subsequently sterile ice chips of equal size were added to each sample to initiate nucleation. The temperature was then lowered to -10 °C at 0.01 °C sec^-1^. Temperature measurement points were set with the software for each leaf ∼1 cm from the wound and recorded as freezing progressed. Data was analysed using Logger Pro (version 3.14) (Vernier Software and Technology). The assays were performed in triplicate.

Assessment for whole plant freeze survival was modified from previous methods (Colton-Gagnon et al., 2014; Bredow et al., 2016; Mayer et al., 2020). *Brachypodium* seeds were sterilised and sown with 10 seeds per line and condition, and evenly spaced apart in 4” x 6” x 3” pots, as described in plant growth conditions. Seeds were then stratified in darkness at 4 °C for 4 days to synchronise germination. Seedlings were grown for two weeks at standard, previously described conditions. The pots were then transferred to another climate-controlled chamber (Econair GC-20; Ecological Chambers Inc.) where plants were exposed to -1 °C for 12 h before the lights were turned off and temperature was ramped down at a rate of 1 °C per hour until -8 °C, and after which the temperature was returned to 4 °C and the lighting resumed (∼150 μmol m^−2^s^−1^) for 24 h. After recovery, plants were returned to standard conditions for 2 weeks, survival was recorded, and images were captured. Survival assays were repeated in triplicate.

Infections at high subzero temperatures were assayed using liquid cultures of *P. syringae* pv. *syringae* B728A, a pathovar with ice nucleation activity (Feil et al., 2005). The bacteria were cultured while shaking at 28 °C to OD_600_ = 0.6-1.0. They were then placed at 4 °C for two days while shaking to increase INP production and resuspended in 10 mM MgCl_2_ and diluted to an OD_600_ of 0.2, corresponding to an approximate concentration of 1×10^8^ colony forming units (CFU) mL^-1^. Simultaneously, three-week-old plants were CA at 4 °C for two days. Extra care was taken to ensure plants and cultures remained separated. Leaves of equal size and length were aseptically removed and the wound was dipped in the bacterial cultures. Leaves were then incubated at -3 °C in an enclosed temperature regulated chamber for 12 h, assessed for evidence of disease including water soaking and cell death and then allowed to recover at 4 °C in the dark with reassessment at 24 h, 48 h, and one week. Preliminary experiments involved plants grown at standard conditions for three weeks with NA and CA plants sprayed with culture as well as cut leaves exposed to wound dipping as described. Whole plants and leaf tissue preliminary infections were carried out at -3 °C and recovered as described with separate plants and leaf tissue maintained at standard *Brachypodium* conditions following infection as controls. All assays were repeated a minimum of three times.

## 3 Results

### 3.1 *BdIRI* Gene Analysis

The number of predicted full-length *IRI* genes in *Brachypodium* database has changed as genome assembly quality and annotation tools have improved. Based on the v3 RefSeq annotation (GCF_000005505.3), *Brachypodium* is predicted to encode six full-length *IRI* genes (*BdIRI1, BdIRI3, BdIRI4, BdIRI5, BdIRI6* and *BdIRI7*) and one gene (*BdIRI2*) encoding a truncated protein with an intact LRR domain but with no AFP domain. Alignments of all amino acid sequences corresponding to *BdIRI1-7* using Clustal Omega from the v3 assembly showed annotated apoplast localization signal sequences, LRR motifs of LxxL where x represents a non-conserved residue, putative protease hydrolysis sites, as well as AFPs (except for *BdIRI2*) with ice-binding motifs of NxVxG/NxVxxG where x represents an outward-facing residue of the beta-barrel structure (Figure S1). The genes are found in three gene clusters on chromosome five (Figure S2). The organisation shows an adjacent position of two to three genes, with transcription in the same direction within each cluster, and flanked by distinct genes including some that may be involved in epigenetic regulation. It is possible that adjacent *BdIRI* loci are similarly spatio-temporally or developmentally regulated. For example, *BdIRI3* and *BdIRI4* are adjacent in the genome, and peptides corresponding to the AFP domain of both of these genes were identified after mass spectrometry of CA *Brachypodium* leaves (Bredow et al., 2016).

To obtain insight into the regulation of the *Bd*AFP genes to better devise a knockdown strategy that targeted AFP expression only after exposure of the plant to low temperatures, sequences upstream of the translational start site of the *BdIRI*s, as well as the promoter region from a known cold-inducible gene from rice, pr*Os*MYBR1R35 (Figure 1; Li et al., 2017), were examined for putative *cis*-acting regulatory elements (CAREs) using PlantCARE (Figure 1; Table S1). All sequences were found to contain *cis*-elements including canonical promoters or enhancers such as TATA-boxes and CAAT-boxes. Cold response-related and drought-resistant (CRT/DRE) core motifs (CCGAC), which are *cis*-elements involved in low-temperature stress responses, were associated with all the promoters. Additional motifs associated with stress and low-temperatures including various abscisic acid-responsive elements (ABREs), low-temperature response elements (LTREs), inducer of cold or C-repeat binding factor expression 1 CBF expression 1 (ICEr1), drought response elements (DREs), and the WRKY stress transcription factors recognition W-box motifs were all found, and these have been annotated (Figure 1). The cold-inducible rice promoter shared with the *Bd*AFP promoters multiple CRT/DRE motifs in addition to other elements involved in low temperature regulation and thus it was surmised to be suitable to drive expression of the inhibitory miRNA.

**Figure 1.**
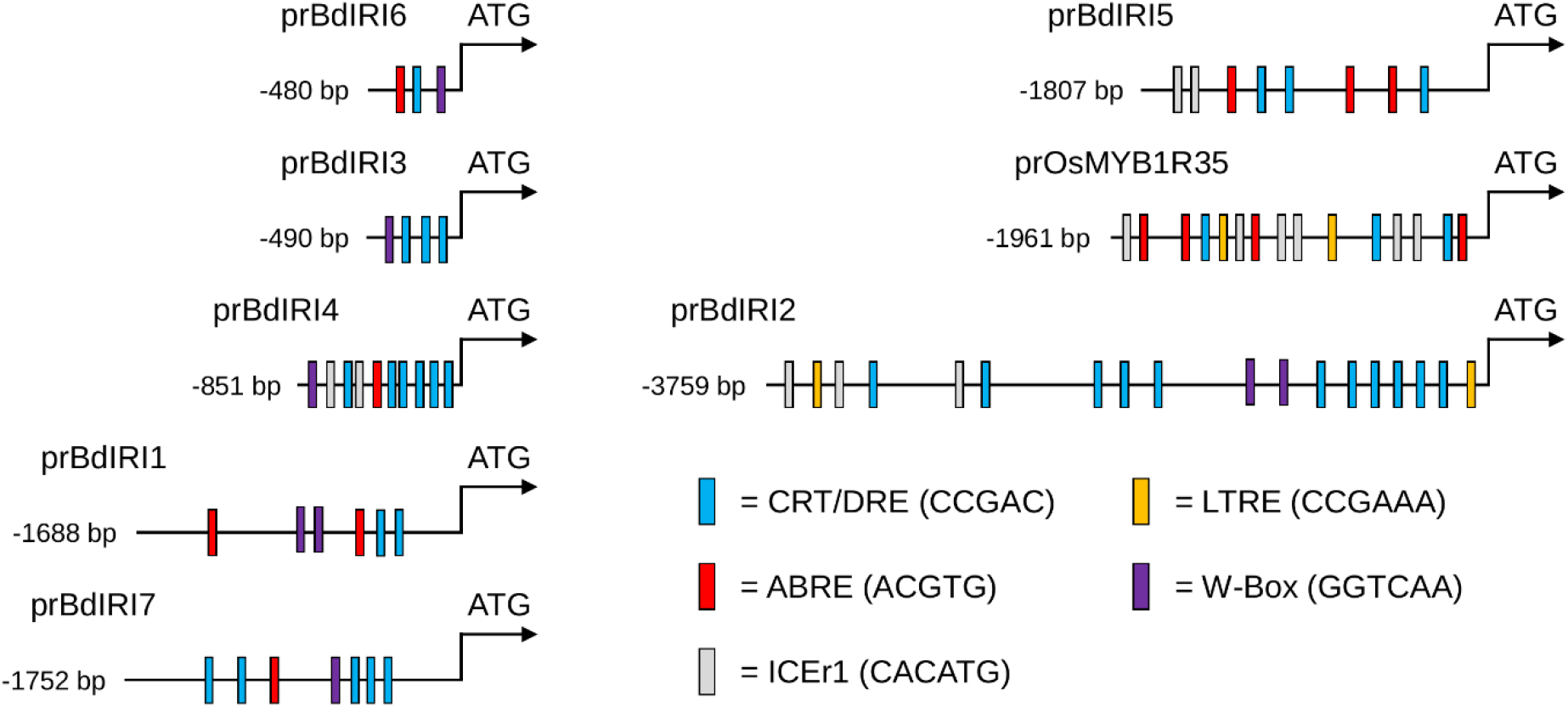
Illustrations of putative promoter regions 5’ of the ATG start codon in the *Brachypodium distachyon* AFP genes *BdIRI1-7* and the known cold-regulated promoter sequence of the rice, *Oryza sativa*, gene *OsMYB1R35*. The analysis extended until the stop codon of the nearest upstream gene. *Cis*-regulatory elements are annotated with strand positions shown relative to the sequence encoding the ATG start. Coloured boxes correspond to canonical cold response related and drought resistant element motifs (CRT/DRE; blue), the stress hormone, abscisic acid-responsive elements (ABREs; red), the cold response pathway inducer of C-repeat binding factor expression 1 (ICEr1; grey), low-temperature response elements (LTREs; yellow), and the WRKY stress transcription factors recognition W-box motifs (purple).

Plant gene expression can be regulated by miRNAs both transcriptionally (Yang et al., 2019) and post-transcriptionally (Jones-Rhoades et al., 2006; Mallory and Vaucheret, 2006; Bertolini et al., 2013; Zhang, 2015), with *Brachypodium* known to transcribe miRNAs as part of its cold-stress response (Zhang et al., 2009). Among monocots, *Brachypodium* and rice have the highest number of annotated miRNAs with 525 and 713 (*O. sativa*) in the miRBase database (Release 21). Again, to inform the knockdown strategy, all 7 *BdIRI* genes, as well as the rice promoter, pr*Os*MYBR1R35, were analysed for potential miRNA binding sites. Within the *BdIRI* coding regions, the majority of miRNAs (14/28), belong to the bdi-miR395 family which have homologs in rice as well as *Arabidopsis* (Zhang et al., 2009). This miRNA family is stress-regulated with members known to target disease resistant proteins (Jones-Rhoades et al., 2004; Fujii et al., 2005; Baev et al., 2011; Lv et al., 2016). Other miRNAs from the miR169 family had 6 putative targets in *BdIRI2* and *BdIRI5* and are predicted to be involved in the plant oxidative stress response (Lv et al., 2016). Likewise, other miRNAs are predicted to regulate multiple *BdIRI*s including bdi-miR7717a-5p on *BdIRI2* and *BdIRI3* and bdi-miR5055 on *BdIRI2, BdIRI3, BdIRI5*, and *BdIRI6*, suggesting that *BdIRI*s can be regulated by common miRNAs, in addition to other miRNAs that target individual *BdIRI* transcripts. For example, *BdIRI4, -3* and *-1* are clustered on the chromosome but only *Bd*AFP isoforms 3 and 4 were found after mass spectrophotometric analysis of leaves (Bredow et al. 2016), suggesting that the lack of the *Bd*AFP isoform 1 in that tissue may be due to miRNA regulation by bdi-miR159b-3p.2 with a binding site in the *BdIRI1* transcript but not in the other *BdIRI* transcripts. In total, the algorithm indicated that all of the *BdIRI*s had at least one predicted miRNA target in the corresponding transcript and all but *BdIRI4* showed miRNA binding sites upstream of the coding region (Table S2). Notably the 1961 bp rice promoter region also showed multiple *Brachypodium* miRNA target sites, as did sequences corresponding to *BdIRI1, -2, -5*, and *-7* gene promoters (Table S2). The observation that the low temperature-regulated promoter regions from both species shared some *Brachypodium* miRNA target sites (*e*.*g*. miR1583 and miR5174d), again suggested that the choice of this rice promoter to drive expression of the miRNA in *BdIRI* knockdown lines was likely appropriate.

### 3.2 Promoter Function and Developmental Phenotypes

In an attempt to curtail pleiotropic effects that might have been due to the constitutive expression of the miRNA in previous knockdown constructs (Bredow et al., 2016), CaMV 35S was substituted with the cold-induced rice promoter. After construction of the plasmids, *Brachypodium* was successfully transformed using a seed cut method (modified from Fursova et al., 2012), which from a total of 150 seeds, yielded three and two PCR-positive transformants for the miRNA and eGFP constructs, respectively. The rapid genotyping method (Ben-Amar et al., 2017) proved effective and overall, 65% of the recovered T_1_ generation were resistant to the hygromycin-selective media.

Once homozygous lines were selected, the ability of the heterologous rice promoter to direct transcription was assessed using western blots with transgenic control *Brachypodium* bearing the pr*OsMYB1R35*:eGFP construct (Figure S3). No bands corresponding to GFP were detected in NA or CA wild type extracts, nor in NA pr*OsMYB1R35*:eGFP leaves. However,, CA pr*OsMYB1R35*:eGFP extracts showed a band at 26 kDa that co-migrated with purified GFP. This demonstrates the successful expression of a marker protein in *Brachypodium*, driven by the *OsMYB1R35* promoter from *O. sativa*.

Plants bearing the rice promoter ligated to the miRNA sequence appeared to develop similarly to wild type and showed normal phenotypes with respect to height and seed production (Figure S4). Two homozygous lines were identified and designated prOmiRBdIRI-1e and prOmiRBdIRI-3c. The prOmiRBdIRI-1e and prOmiRBdIRI-3c lines were 25.8±4.07 cm and 21.86±4.37 cm, and set 106.2±29.9 and 99.5±32.6 seeds, respectively, which was not significantly different from wild type at 23.1±4.1 cm and 101.4±27.7 seeds (unpaired *t*-tests; assessed at 12 weeks using three independent growth trials with at least 15 plants per trial; Figure 2). Germination rate was also the same at 90.4±10.9%, 90.0±13.1%, and 91.25±11.3%, for wild type, prOmiRBdIRI-1e, and prOmiRBdIRI-3c lines, respectively (8 independent growth trials using at least 10 seeds per trial).

**Figure 2.**
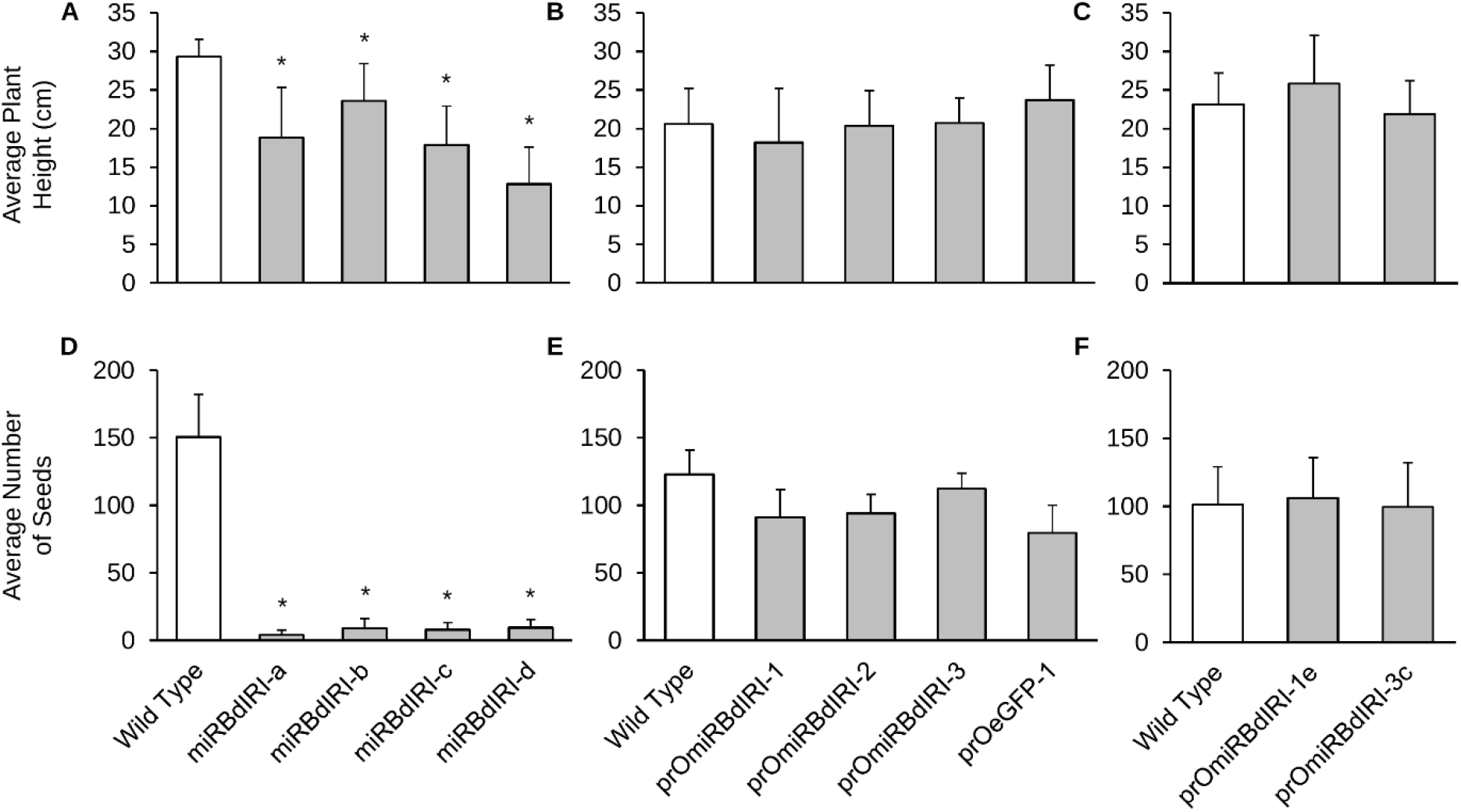
The average height and number of seeds per plant, in *Brachypodium* Bd21 wild type plants and plants from transgenic lines. **(A**,**D)** Homozygous transgenic lines employing the CaMV 35S promoter ligated to the miRNA sequence to generate constitutively expressed *BdIRI* knockdown plants (data taken from Bredow et al.,2016). **(B**,**E)** Heterozygous temporal knockdown lines employing the *OsMYB1R35* promoter, and **(C**,**F)** homozygous temporal knockdown lines. The data in B, C, E, and F and was compiled at 12 weeks from three independent growth trials using at least 15 plants per trial for each knockdown line and wild type. Asterisks indicate significant differences compared to wild type (unpaired *t*-test, *p* < 5 × 10^−9^).

### 3.3 Antifreeze Activity and Freeze Resistance

The ability to knockdown AFP expression was tested in “splat” assays to visualise IRI activity in crude extracts or apoplast samples (Figure 3, Figure S7). CA knockdown prOmiBdIRI-1e and prOmiRBdIRI-3c lines all showed reduced AFP activity compared to wild type CA plants and were similar to assays of NA wild type and NA transgenic lines. A dilution series used to estimate the difference in activity showed that at 0.01 mg mL^-1^, CA knockdown lines and NA wild type show larger ice crystals at the conclusion of the annealing period compared to samples from CA wild type and CA prOmiReGFP. Thus, CA appears to regulate the rice promoter to drive the miRNA to attenuate the expression of the AFP gene products, but it is possible that endogenous *BdIRI* promoters may have some very low levels of expression under NA conditions. As well, although the two lines both show attenuation of AFP activity, there appeared to be minor differences in expression, undoubtedly due to position effects, as has been shown in other transgene insertions (e.g. Simón-Mateo and Garcia, 2006; Duan et al., 2008; Liu et al., 2017).

**Figure 3.**
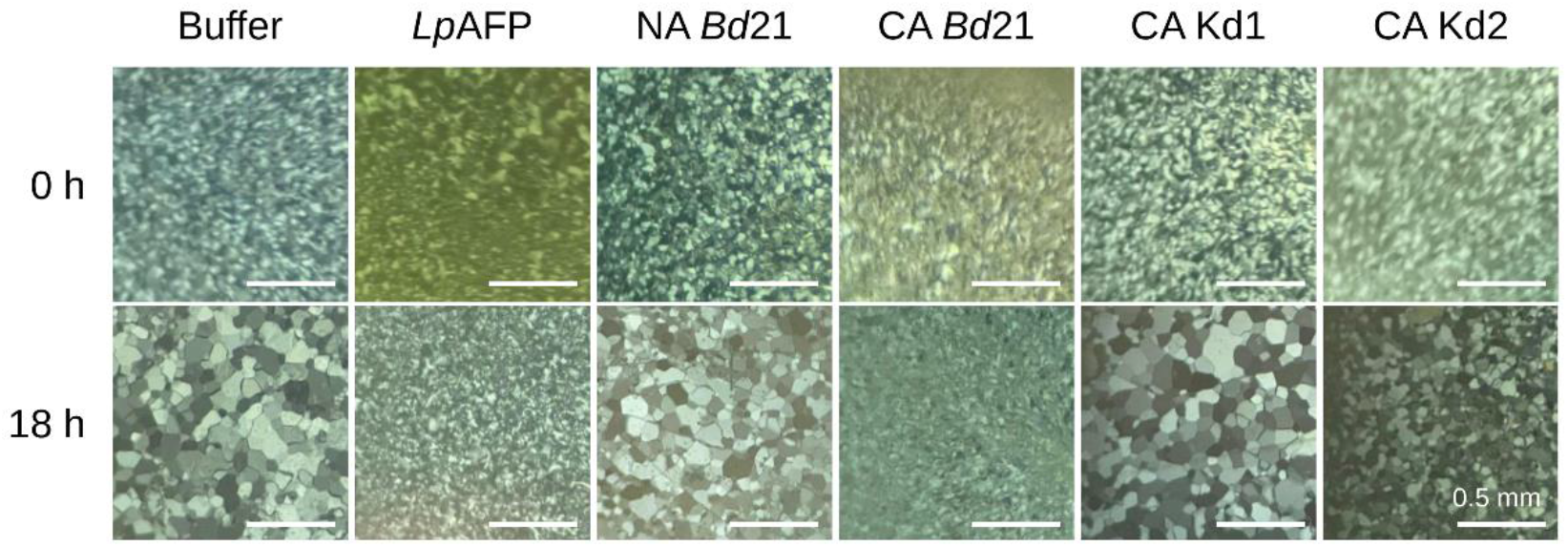
Ice recrystallization inhibition “splat” assays of apoplast extracts using non-acclimated (NA) and cold-acclimated (CA) *Brachypodium distachyon* Bd21 wild type and temporal antifreeze protein knockdown lines prOmiRBdIRI-1e (Kd1) and prOmiRBdIRI-3c (Kd2). Samples were annealed at -6 °C for 18 h at a standardised concentration of 0.01 mg mL^-1^. Buffer and recombinant purified rye grass (*Lolium perenne* AFP; *Lp*AFP) controls are also shown. Assays were performed in triplicate with similar results, and representative images are shown. Scale bars represent 0.5 mm. Splat assays performed using CA prOmiReGFP plants resulted in results similar to those obtained using wild type Bd21, indicating that plasmid presence and seed transformation did not affect AFP activity (not shown).

TH was significantly increased to 0.05 °C in wild type tissue extracts from CA plants compared to being virtually undetectable in NA samples (Table 1; Figure S5). Plant AFPs have TH levels that are typically low with ice-purified *Bd*AFPs previously reported at 0.08 °C (Bredow et al., 2016). CA transgenic prOmiRBdIRI-1e and prOmiRBdIRI-3c had TH activities that were significantly reduced (77% and 74%) compared to the levels shown by CA wild type. Strikingly, these low-temperature induced transgenic knockdowns showed levels of TH at 0.013 °C and 0.012 °C, comparable to previously reported constitutive knockdowns expressing the same miRNA, at a TH range of 0.009-0.034 °C (Table 1 and Bredow et al., 2016), confirming the successful silencing of *Bd*AFP activity even when the miRNA was driven by the temporal rice promoter.

**Table 1.** Thermal hysteresis (TH) readings done using crude protein extracts from leaf tissue lysates on cold-acclimated (CA) and non-acclimated (NA) Bd21 wild type and temporal cold-induced antifreeze protein knockdown lines prOmiRBdIRI-1e and prOmiRBdIRI-3c. Samples were tested at 40 mg mL^-1^ of total protein concentrated from crude cell extracts. Readings were captured using a nanoliter osmometer. Assays were performed in triplicate and values shown are the average of three replicates with standard deviation.

| Sample | TH (°C) |
| --- | --- |
| Buffer | 0 |
| Wild Type, Bd21 (CA) | 0.050±0.019 |
| Wild Type, Bd21 (NA) | 0.005±0.003 |
| prOmiRBdIRI-1e (CA) | 0.013±0.002 |
| prOmiRBdIRI-1e (NA) | 0.003±0.003 |
| prOmiRBdIRI-3c (CA) | 0.012±0.011 |
| prOmiRBdIRI-3c (NA) | 0.002±0.002 |

*Brachypodium* AFPs shape ice into hexagon crystals followed by a smooth, irregular, flower-shaped burst (Figure S6). As would be expected, samples from CA wild type plants showed obvious ice shaping, but some minimal shaping still occurred in NA wild type and knockdown samples (Figure S7), consistent with the IRI results. Ice shaping in the CA knockdowns appeared to occur on the primary prism plane, favouring slight shaping along the *a*-axis before weakly bursting (Figure S7). In comparison, the CA wild type showed initial adsorption affinity for the *a*-axis primary prism plane quickly followed by adsorption and shaping on the *c*-axis basal plane, forming stunted hexagonal bipyramidal forms with strong bursts when the freezing point was exceeded. Rounded *Bd*AFP-mediated ice burst morphology presumably would help protect membranes by preventing the growth of large, sharp ice crystals as is seen in some freeze-avoiding organisms (Bar Dolev et al., 2016). Such membrane protection can be quantitatively assessed by electrolyte leakage assays. The CA transgenic leaves showed a significant increase (∼30%) in electrolyte leakage (*p* < 0.05, one-way ANOVA) at -10 °C when compared to CA wild type (Figure 4). At higher freezing temperatures of - 6 °C, electrolyte leakage was variably increased in the CA knockdown lines relative to CA wild type leaves (not shown). In all experiments, leaves from plants maintained at 4 °C showed relatively little electrolyte leakage independent of the genotype, as expected.

**Figure 4.**
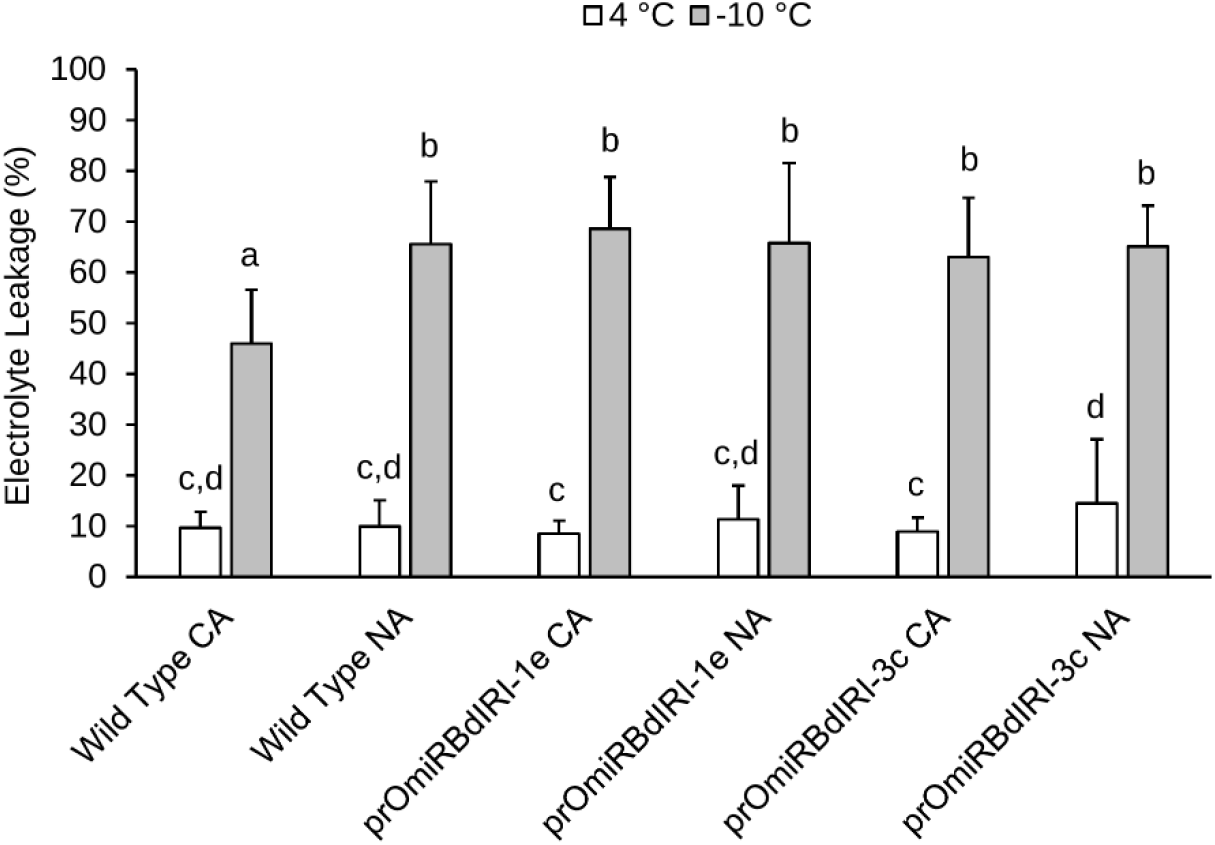
Electrolyte leakage assays performed on non-acclimated (NA) and cold-acclimated (CA) wild type *Brachypodium distachyon* Bd21 and two homozygous temporal antifreeze protein knockdown lines (prOmiRBdIRI-1e and prOmiRBdIRI-3c). Control leaves were maintained at 4 °C (white bars) while experimental samples were incubated at -10 °C (grey bars) as indicated. Electrolyte leakage was measured as a percentage of electrolytes released after the freeze protocol as a ratio of the total released electrolytes after autoclaving, based on the total leaf mass in the sample (see Methods). Letters represent statistically significant groups following one-way ANOVA with post-hoc Tukey multiple test correction (*p* < 0.05). Error bars represent the standard deviation of the mean and assays were performed in triplicate (n = 10).

When entire plants were frozen to -8 °C, 47% of CA wild type survived, significantly more (*p* < 0.01, unpaired *t*-test) compared to 20% of CA prOmiRBdIRI-1e plants and none of the CA prOmiRBdIRI-3c plants (Figure 5). In addition, none of the NA plants survived, independent of genotype. To further assess ice propagation and freezing patterns of the temporal knockdown compared to wild type plants, infrared thermography was used as encouraged by previous observations of ice nucleation and propagation in various species (Wisniewski et al., 1997; Lutze et al., 1998; Ball et al., 2002; Sekozawa et al., 2004; Wisniewski et al., 2015). Images of CA and NA freezing leaf tissue from wild type compared to knockdown lines as the temperature was reduced to - 10 °C were distinct, suggesting that AFPs can slow the propagation of ice through leaf tissues, and were consistent with the electrolyte leakage assays (Figure 6). Thermographs further supported the conclusion that ice propagation at subzero temperatures was more rapid in leaves from AFP knockdown lines. Temperature readings collected concurrently with thermographs on leaves show CA wild type leaves are 1-2 °C warmer than the NA wild type and CA knockdowns, consistent with the known ∼2 °C freezing point depression of INPs mediated by *Bd*AFPs with a divergence around -2 °C where INPs nucleate ice (Figure S8).

**Figure 5.**
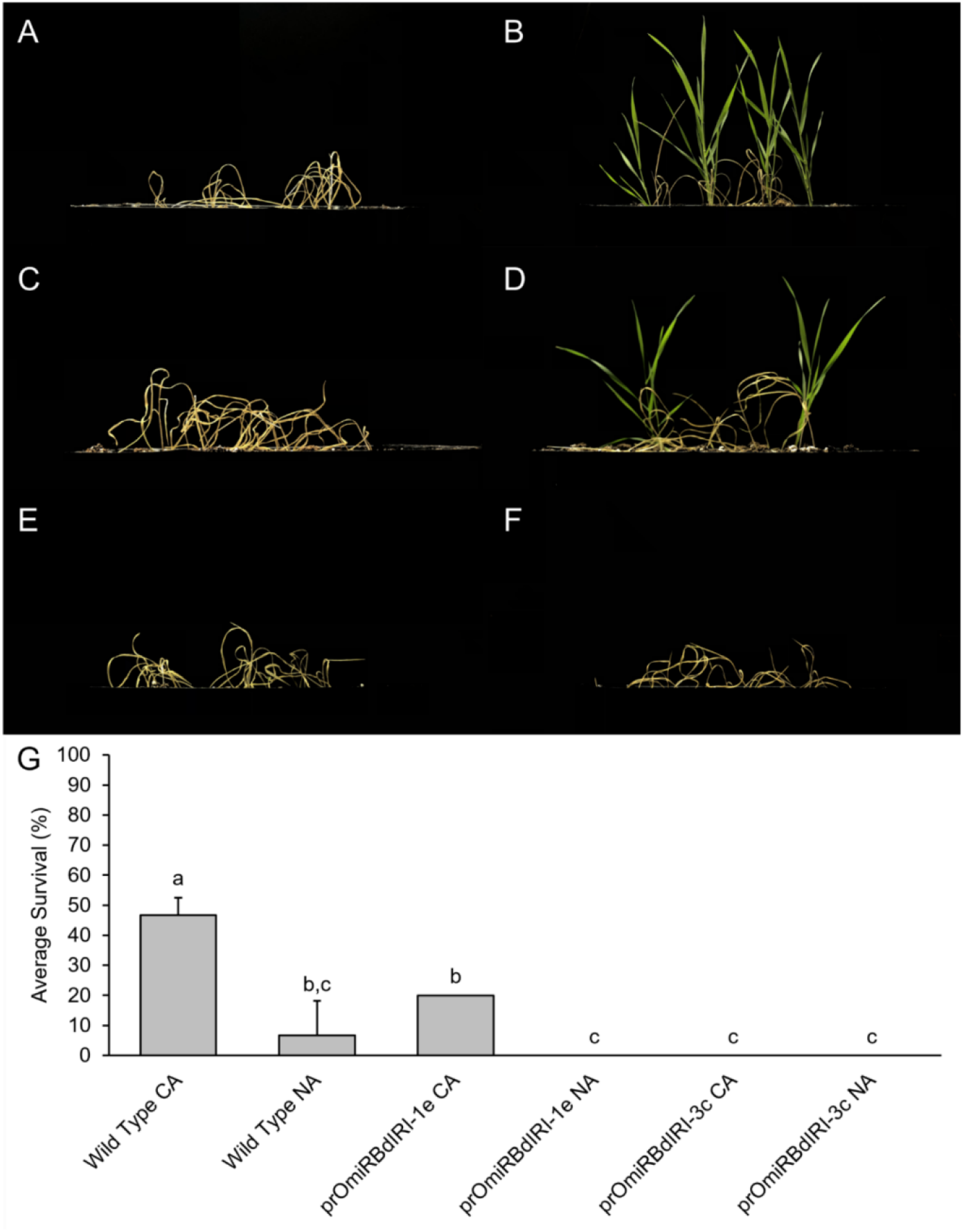
Whole plant freezing survival assay of non-acclimated (NA) and cold-acclimated (CA) wild type *Brachypodium distachyon* Bd21 and two homozygous knockdown lines (prOmiRBdIRI-1e and prOmiRBdIRI-3c). **(A)** NA Bd21 wild type. **(B)** CA Bd21 wild type. **(C)** NA prOmiRBdIRI-1e. **(D)** CA prOmiRBdIRI-1e. **(E)** NA prOmiRBdIRI-3c. **(F)** CA prOmiRBdIRI-3c. **(G)** Values represent survival and are the average of three replicates, with error bars showing the standard deviation of the mean. Letters represent statistically significant groups following one-way ANOVA with post-hoc Tukey multiple test correction (*p* < 0.01). Two-week-old plants were frozen at a rate of 1 °C h^-1^ to -8 °C in a temperature-controlled chamber in the dark following misting with sterile water to initiate freezing. Plants were allowed to recover for two days at 4 °C with no light and returned to standard conditions for 7 days before images (A-F) were captured. Assays were performed in triplicate (n = 10).

**Figure 6.**
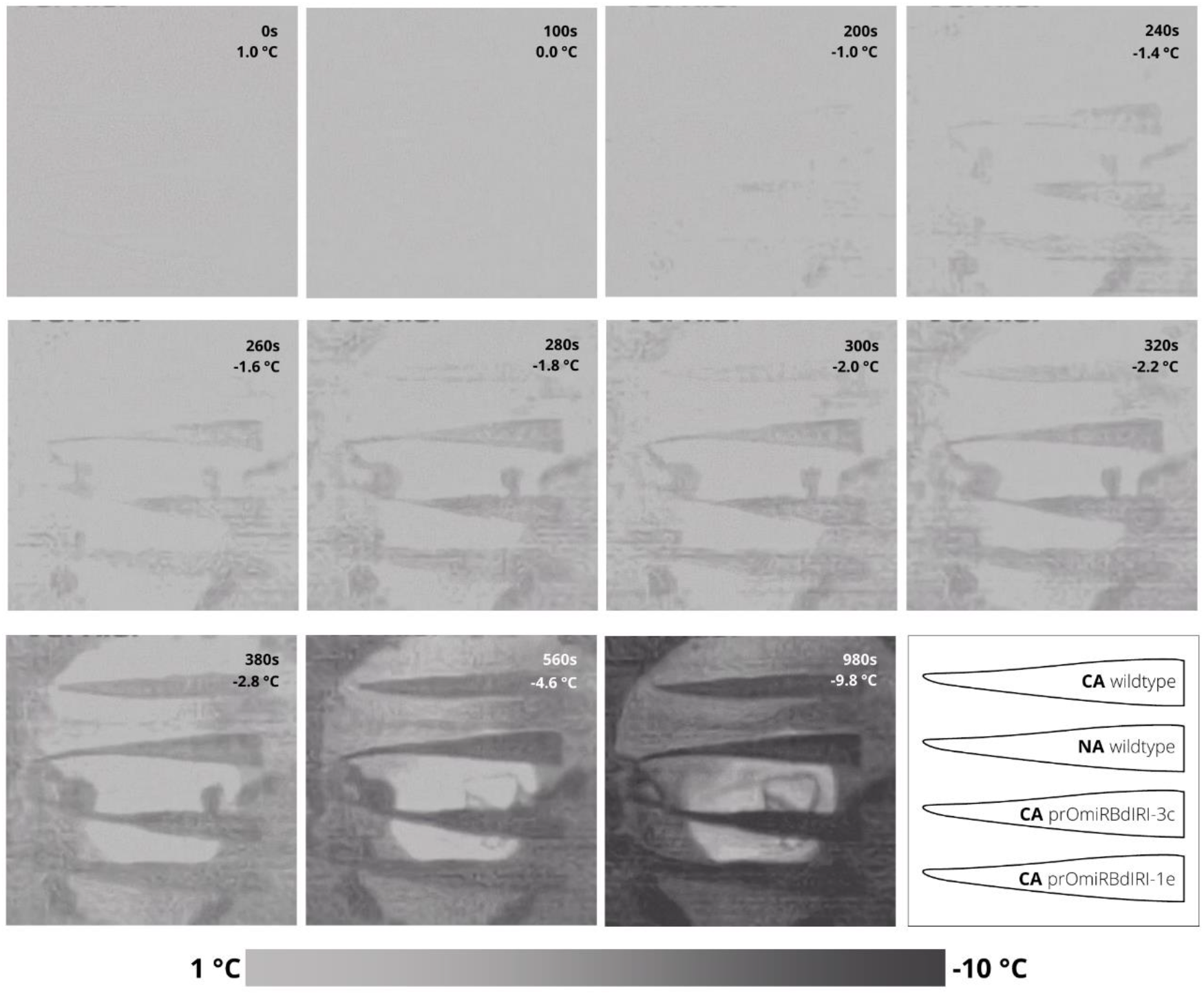
Thermographs of excised leaf tissue of wild type cold-acclimated (CA), wild type non-acclimated (NA), prOmiRBdIRI-1e CA, and prOmiRBdIRI-3c CA, from top to bottom, respectively. Leaves were equilibrated at 1 °C and frozen to -10 °C. Ice propagation was nucleated by an ice chip placed in 10 μL of water on the excision wound. Time stamps and temperatures are indicated. A diagram of the leaf samples is shown in bottom right with a temperature scale shown at the bottom.

Ice nucleation under natural conditions is invariably initiated by INA+ bacteria, including strains of *P. syringae*, but as noted, *Brachypodium* AFPs can attenuate INPs *in vitro* (Bredow et al., 2018). To measure the impact of *Bd*AFPs on pathogen infection, wild type and a knockdown line were exposed to *P. syringae* pv. *syringae* B728A, a pathovar with INPs (Feil et al., 2005). Leaves from NA wild type and CA knockdown lines treated with the bacterial culture and placed at -3 °C, a temperature below which this pathovar nucleates ice (Figure S9A), displayed disease-like symptoms including water soaking and cell death, while infected plants that were not subjected to freezing temperatures displayed non-freeze-associated infection symptoms 12 h post infection (Figure S9B-S9E). In contrast, CA wild type leaves displayed little evidence of tissue damage (Figure S10). After 24-48 h post infection, leaves from plants known to have little or no AFP activity were shrivelled and dry compared to CA wild type leaves, likely due to cell lysis associated with ice nucleation and growth. These same leaves showed more disease symptoms and cell death one week after pathovar exposure compared to CA wild type controls. Although it is not surprising that treated CA wild type leaves showed symptoms of disease considering the bacterial titre used, they appeared to have greater resistance to dehydration compared to the leaves with low AFP activity. This qualitative assay suggests that AFPs could ameliorate the impact of pathogens with INA activity.

## 4 Discussion

### 4.1 Heterologous Promoter-Driven Expression

The decision around the selection of an appropriate promoter to drive transcription in transgenic organisms is frequently challenging. Regulatory sequences including enhancers associated with the promoter can dictate tissue and developmental expression and thus transcription of a coding sequence of interest can be finely tuned particularly in model organisms where sequence libraries of promoters and enhancers are available, such as the rich resources available for *Drosophila* geneticists. For grasses, genetic resources are scarcer and in the absence of annotated promoter sequences, constitutive promoters such as CaMV 35S are used. However, their employment can risk inappropriate expression, epigenetic silencing, and even suboptimal growth (Estrada-Melo et al., 2015; Rajeevkumar et al., 2015; Amack and Antunes, 2020). Thus, the short stature, low germination rate, and the near-sterility in transgenic plants generated using the CamV 35S promoter ligated to a sequence for a miRNA that targets the translation of *BdIRI* gene products was unfortunate but not unexpected. Previous empty vector control plants did not present pleiotropic effects (Bredow et al., 2016), and thus these detrimental characteristics could be associated with the constitutive presence of the miRNA, and either due to the attenuation of AFPs and/or LRRs perhaps needed during stem elongation and seed formation, or simply the expression of these unregulated miRNAs recruiting polymerases or polysomes and thus interfering with proper development.

As an alternative to constitutive promoters, synthetic promoters consisting of core promoters and combinations of enhancer sequences for regulatory elements, sometimes from heterologous species, can be constructed with uncertain outcomes (Mohan et al., 2017; Ali and Kim, 2019). Again, as another somewhat risky strategy, an entire promoter sequence from evolutionarily related plants with documented expression profiles similar to the target genes in a second species can be selected and may allow for a greater degree of control including spatio-temporal expression (Dutt et al., 2014). Likewise, heterologous promoters may be free from endogenous signalling that could lead to undesired effects if using a native host promoter (Napoli et al., 1990). For the *Bd*AFPs that are known to be produced within two days of CA, the cold-regulated promoter from *O. sativa* was an attractive choice even though we were unaware if it had been previously used heterologously. In rice it is induced at 4 °C, the CA temperature of *Brachypodium*, and the directed rice transcripts increase continuously for 48 h (Li et al., 2017). Here we show that the use of this rice promoter indeed functions in *Brachypodium* and there were no obvious detrimental phenotypes associated with its use, either in the transgenics bearing the miRNA sequence or in the eGFP expression plants (Figure 2).

Non-coding RNAs including miRNAs are part of a complex signalling network having roles in developmental regulation and stress response with multiple targets for single miRNAs (Chen, 2009; Peter, 2010; Wu et al., 2010; Budak and Akpinar, 2015; Liu et al., 2017; Waititu et al., 2020). Complementary sequence targets include gene transcripts that can be hydrolysed, translationally inhibited, or can localise to the nucleus and target promoter sequences (Jones-Rhoades et al., 2006; Li et al., 2013; Yang et al., 2019). Their frequent roles in the regulation of stress responses are of interest with regards to genes induced after CA. The *Brachypodium* Poaceae lineage diverged from rice ∼50 million years ago, with high homology and synteny remaining between *Brachypodium* and rice genomes (Bossolini et al., 2007; Huo et al., 2009; Kumar et al., 2009; IBI, 2010). The identification of *Brachypodium* miRNA predicted target sequences in both the *BdIRI* and the *OsMYB1R35* promoters suggests an evolutionarily conserved low temperature response that encouraged the prospect that the rice promoter would not only be recognized but could be appropriately regulated by any of the identified *Brachypodium* miRNAs with target sites in the rice sequence (Table S1; Table S2).

### 4.2 AFP Knockdowns and Freezing Vulnerability

As noted, the transgenic lines appeared to have no developmental defects and as such the observation that they displayed similar freeze susceptibility as those lines bearing a constitutive promoter driving the miRNA translational interference of AFPs convincingly demonstrates that AFPs do contribute to freeze tolerance. In wild type *Brachypodium*, a short two-day CA period is both necessary and sufficient to prepare the plants for survival to subzero temperatures and is coincident with the appearance of AFPs. These AFPs depress the freezing point of solutions only marginally, but more importantly depress pathogen-mediated INA, shape ice, restrict ice crystal growth as measured by IRI, reduce electrolyte leakage, and ultimately allow whole plant survival even after freezing. Especially striking was the killing of all plants in one line and a second line showing less than half the survival rate compared to wild type controls at -8 °C. All these AFP-related properties were effectively knocked down by the temperature-regulated response of the rice promoter to direct miRNA expression (Figures 3, 4 and 5).

It has been assumed that AFPs are associated with the plasma membrane since reports of a fish AFP bound to model lipid bilayers changed the phase transition and prompted the researchers to recommend the use of fish AFPs to confer low temperature survival to plants, which ultimately did not meet with success (Tomczak et al., 2002; Kenward et al., 1993; Kenward et al., 1999). In plants, the localization of AFPs in the apoplast would suggest limited access to plasma membranes in any case. Indeed, mass spectrophotometry of CA *Brachypodium* plasma membrane proteins did not reveal any *Bd*AFPs (Juurakko et al. 2021b). Nonetheless, AFPs do protect plasma membranes from damage, presumably from uncontrolled ice growth initiated by nucleators in the apoplast with its low solute concentration, and across cell walls into the cells.

The apoplast location of the AFPs allowed their assay in the absence of many other contaminating proteins and yielded clear evidence of the knockdown of IRI activity in the CA transgenic lines with the appearance of large ice crystals after the annealing period (Figure 3). Activity in lysates was consistent but not as visually clear, although individual ice crystals were substantially larger in dilute samples from the knockdowns compared to controls. As judged by western blot analysis of samples from transgenic plants expressing eGFP, the cold-induced rice promoter was not strong in *Brachypodium*. Thus, it is surprising that after a two day induction period, the heterologous sequence-directed miRNA lines showed AFP activities similar to the earlier-reported constitutively expressed miRNA lines with respect to IRI, electrolyte leakage, whole plant freezing survival, and TH (Figures 3, 4, 5, and S5 *vs*. Bredow et al., 2016). This suggests that the single miRNA designed to complement the multiple *BdIRI* transcripts was very effective notwithstanding the heterologous promoter and the miRNA’s less than perfect “match”, underscoring the effectiveness of knockdown by translational attenuation (Bredow et al., 2016).

Structural ice barriers and thermal decoupling controlled by plant architecture and morphology, well known in angiosperms between stems and flowers, may not be as easily applied to grasses (Kuprian et al., 2014; Bertel et al., 2021). No evidence of either were observed in freezing *Brachypodium* leaves analysed by infrared thermography. However, the presence of AFPs in CA wild type tissue was coincident with the obvious retarded advancement of freeze fronts, a lag that was on average 1-2 °C different in leaves with AFPs than without, and coincidentally similar to the ∼2 °C attenuation of ice nucleation activity by *Bd*AFPs (Bredow et al., 2018, Figure S9). There was no inhibition of ice front development observed in NA wild type and CA AFP knockdown lines suggesting that AFPs can delay ice propagation in leaf tissue, consistent with IRI, electrolyte leakage, and whole plant freezing assays. Since mass spectrophotometric analysis has shown that *Bd*AFPs encoded by *BdIRI3* and *4* were detected in leaf tissue (Bredow et al., 2016), at least these two isoforms are associated with leaf ice growth protection. Adsorption of these AFPs to forming ice crystals will keep crystals small and less damaging to adjacent membranes thus limiting leaf damage.

Similar to the attenuation of freezing by AFPs in leaves was the retardation of damage subsequent to infection as seen after exposure of wounded leaves to *P. syringae* pv. *syringae* B728A. This pathovar can nucleate ice at high subzero temperatures presumably as a means to destroy tissues and access nutrients (Feil et al., 2005). It is known that *Bd*AFPs inhibit the activity of the *P. syringae* INP *in vitro* (Bredow et al. 2018) and we speculate that a physical interaction of the INP and *Bd*AFPs is sufficient to impact bacterial fitness, such that the progress of infection was slowed down. This effect was seen as early as 12-48 h after the pathogen challenge, and with the knockdowns showing more cellular death one week post-infection. Thus, it would be of interest in the future to more fully explore the relationship between AFP activity and the infectivity of pathogens bearing INPs.

The cause of the developmental effects seen in constitutive knockdowns of *BdIRI* (Bredow et al., 2016) is unknown. One potential reason may be the miRNA-mediated loss of the LRR isoforms that are hydrolysed in the apoplast from the primary translation products. In general, LRRs function in protein-protein interactions and have roles in growth and development in many organisms, normally in association with various other proteins. As well as LRR-receptor kinases that recognize apoplastic peptides to initiate immune signalling (Fischer et al., 2016) or regulate plant growth and development (He et al., 2018), LRR-extension proteins (LRRX) are bound to cell walls and function in growth as well as pollen tube formation (Zhao et al., 2018). However, to our knowledge, the characterisation and function of plant LRR domains in the absence of adjoining ligands has not been reported and any impact of their knockdown during development is unknown. Instead, the defective phenotypes may have simply arisen due to the constitutive-driven strong miRNA expression non-specifically interfering with gene regulation and polysome loading during critical developmental stages, or alternatively, the possibility that AFPs have an as yet undiscovered role in *Brachypodium* development.

### 4.3 Conclusions and Future Prospects

Taken together, we have shown that substituting a constitutive promoter for a heterologous temporally regulated promoter to direct the expression of a miRNA that targets the *BdIRI* transcripts substantially reduced AFP activity but with no apparent developmental defects. These results highlight the potential of *Brachypodium* AFPs as candidates for the development of freeze-tolerant horticultural crops, if not food crops where there may be public resistance to genetically modified plants. Additional biotechnological and research applications extend outside agriculture and range from additives to prevent ice recrystallization in stored cells and tissues, processed foods, and pharmaceuticals, particularly where infrastructure for flash freezing and extremely low temperature storage is not viable or otherwise unavailable. This research also opens the way to explore *in planta* the effects of AFPs on freezing tolerance and pathogen susceptibility as well as the means by which *BdIRI*s may be regulated through *cis*-regulatory elements and endogenous miRNAs.

## 6 Author Contributions

CLJ conducted all experiments, analysed all data, and produced all figures. CLJ wrote the initial draft of the manuscript and all authors contributed to manuscript revision. MB assisted in the design of some early experiments and conducted preliminary infection experiments. VKW and GCD co-supervised and secured funding.

## 8 Acknowledgments

We acknowledge Dr. Barabara Vanderbeld for her previous published work on the miRNA construct, Dr. Heather Tomalty and Robert Eves (Dr. Peter L. Davies’ lab) for use of the nanoliter osmometer, Jeff Boudreau (Dr. Ian Chin-Sang’s lab) for assistance with the western blots, Kristy Moniz for assisting with plant care, and Ryan Monday for helping to harvest tens of thousands of seeds, phenotyping hundreds of plants, and assisting with plant care.

## 9 Supplementary Material

Supplementary Tables S1 and S2 are available online at: (link here).

---

**Figure S1.**
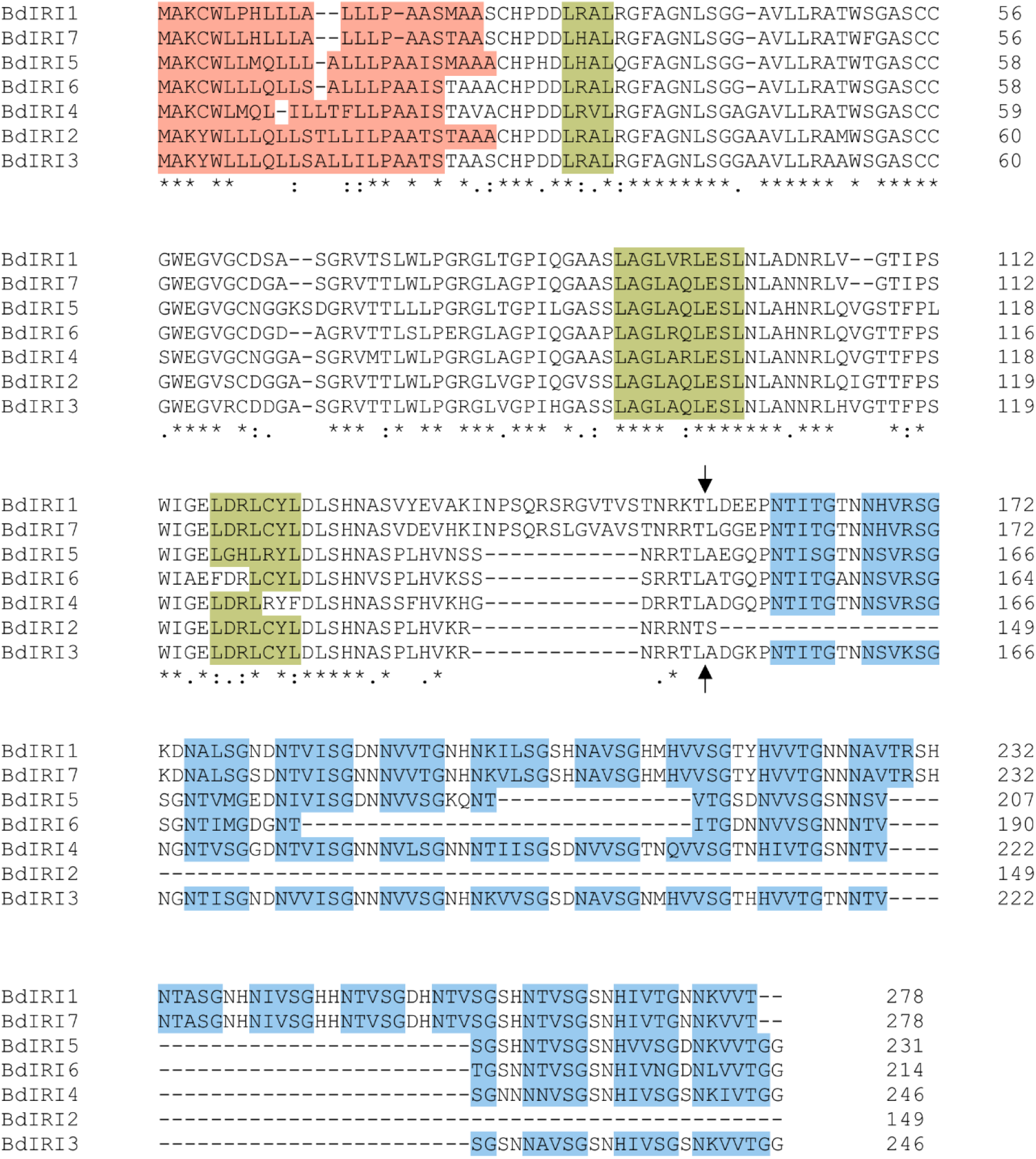
Alignments performed using Clustal Omega (https://www.ebi.ac.uk/Tools/msa/clustalo/) of all 7 *BdIRI* amino acid sequences from the newest assembly (*Brachypodium distachyon* genome v3) containing an annotated apoplast localization signal sequence in red, LRR motifs of LxxL where x represents a non-conserved residue in green, and AFP motifs of NxVxG/NxVxxG where x represents an outward-facing residue of the beta-barrel structure in blue, along with the putative asparagine endopeptidase hydrolytic cleavage sites indicated by black arrows. Asterisks (*) denote fully conserved residues, colons (:) denote conservative substitutions, and periods (.) denote semi-conservative substitutions.

**Figure S2.**
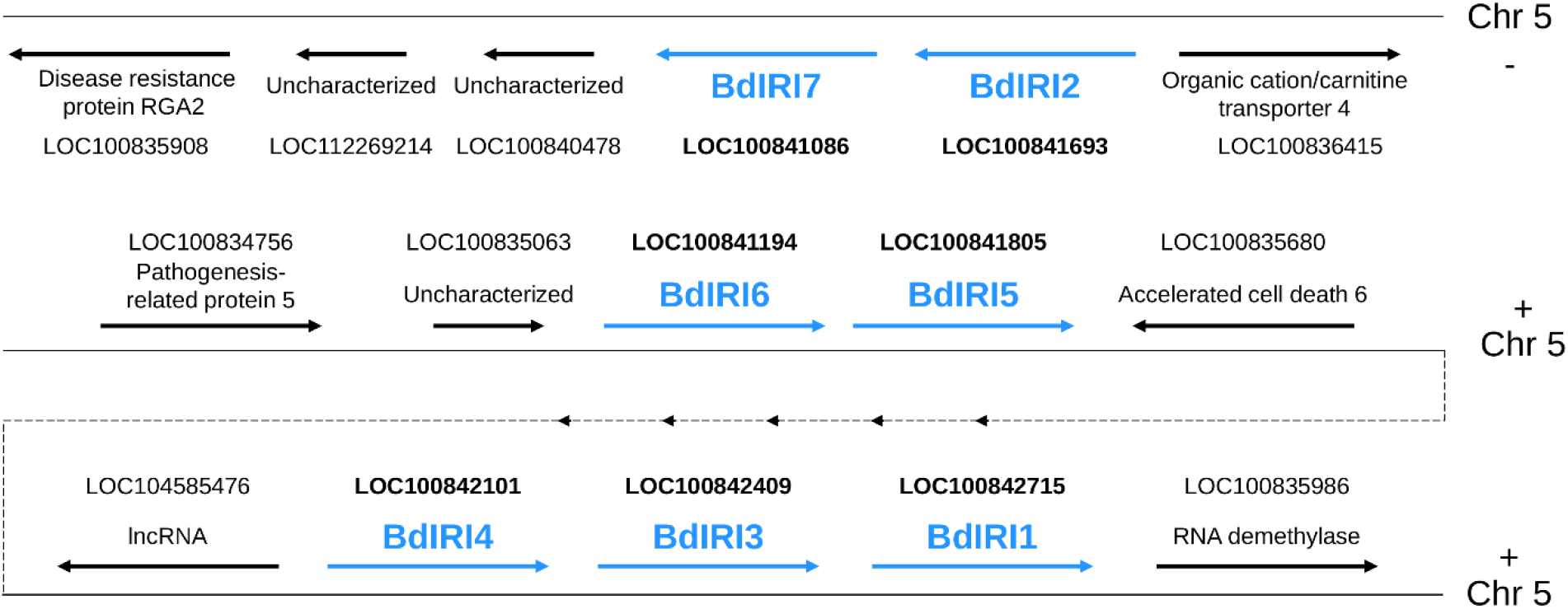
Illustration showing the three clusters and chromosomal positions of the 7 *BdIRI* genes on chromosome five of *Brachypodium* (*Brachypodium distachyon* genome v3). *BdIRI* gene numbers are labelled and strands (+/-) are labelled and are highlighted. Flanking and nearby genes are shown and *BdIRI* genes are highlighted in bold. NCBI RefSeq loci are labelled. Dotted line indicates the direct continuity of the chromosome at this position.

**Figure S3.**
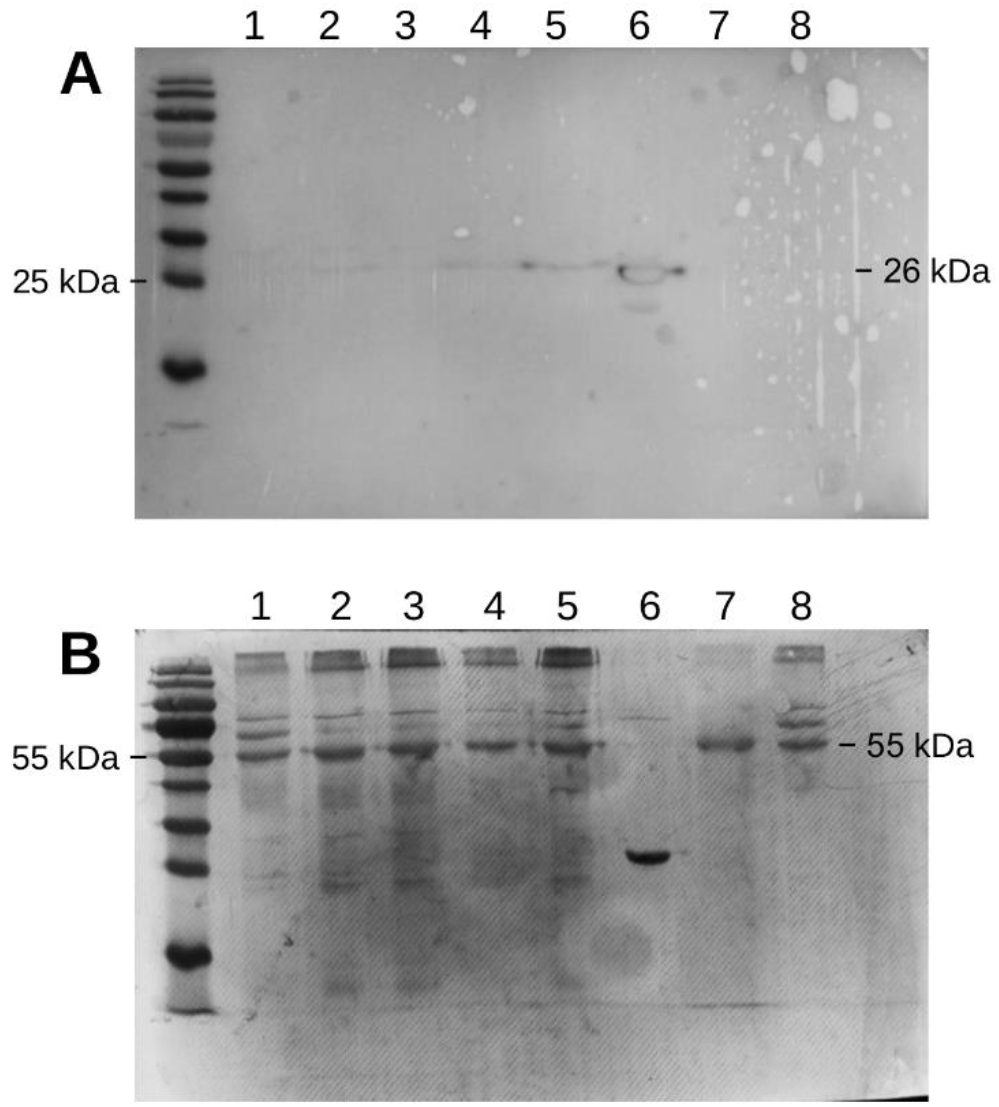
Representative western blot analysis for functional characterization of the rice promoter pr*OsMYB1R35* system in *Brachypodium distachyon* using extracts from non-acclimated (NA) and cold-acclimated (CA) Bd21 wild type and prOeGFP transgenic plants and visualised with antibodies for green fluorescent protein (GFP). **(A)** Lane 1 corresponds to NA prOeGFP, lanes 2-5 to CA prOeGFP from separate individual plants, lane 6 to to purified recombinant GFP used as a positive control, lane 7 to NA wild type, and lane 8 to CA wild type. **(B)** RuBisCO large chain (Rbcl) was used as a loading control with Coomassie Brilliant Blue (CBB) staining. Molecular weights of bands corresponding to eGFP, 26 kDa, and Rbcl, 55 kDa, are labelled. The western blots were performed in triplicate. Note: positive control recombinant eGFP was overloaded and burnt during imaging.

**Figure S4.**
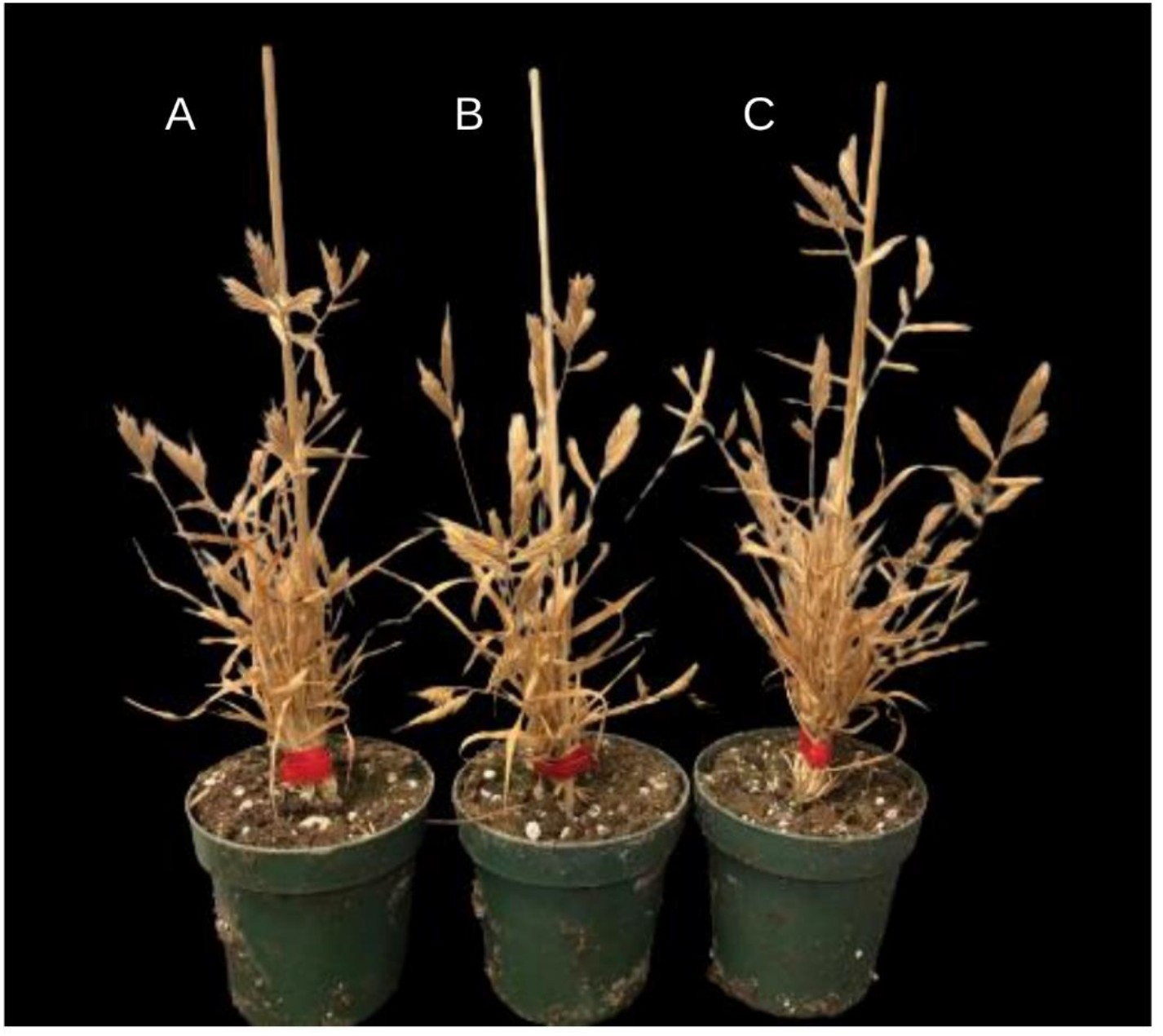
Phenotypes of Bd21 wild type *Brachypodium distachyon* **(A)** plants and two homozygous knockdown lines, prOmiRBdIRI-1e **(B)** and prOmiRBdIRI-3c **(C)**, bearing our temporal AFP knockdown systems. Photos were taken at 12 weeks following the described standard growth conditions with water and fertilizer withdrawn in the final week to allow senesced plants to dry out for seed harvesting.

**Figure S5.**
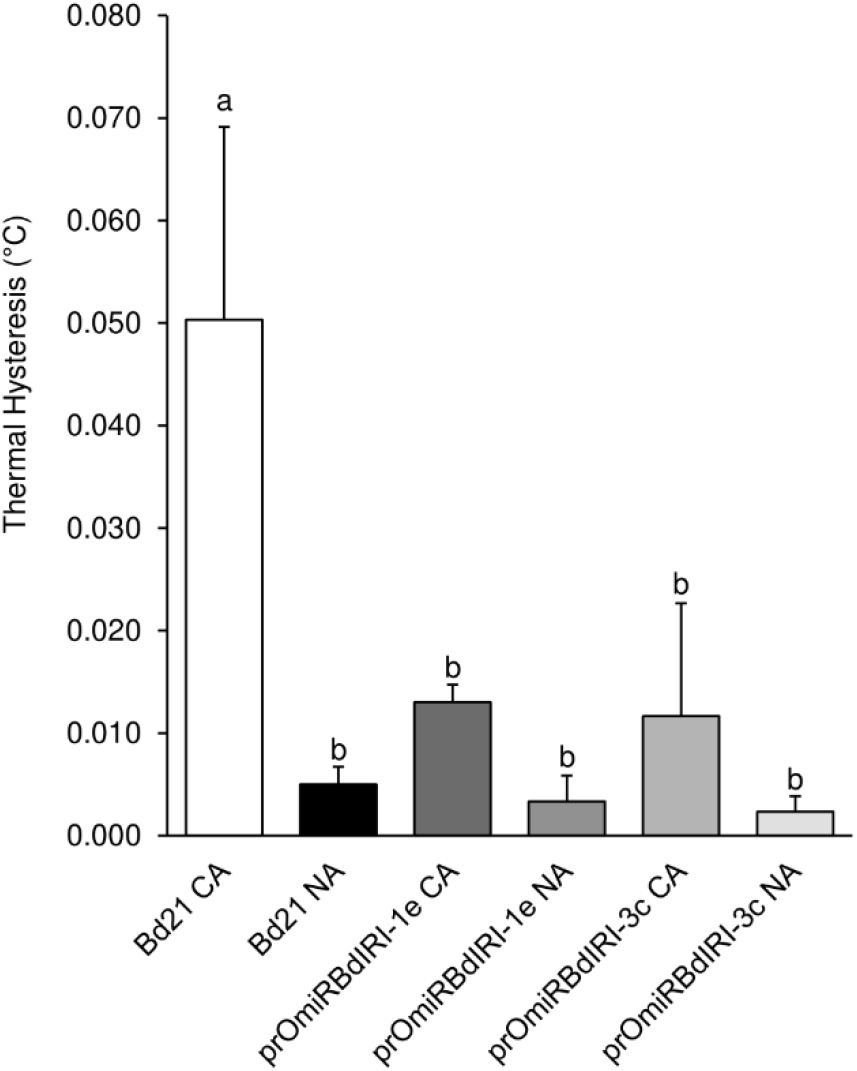
Thermal hysteresis (TH) readings done using crude protein extracts from leaf tissue lysates on non-acclimated (NA) and cold-acclimated (CA) Bd21 wild type *Brachypodium distachyon* and two temporal AFP knockdown lines prOmiRBdIRI-1e and prOmiRBdIRI-3c. Samples were tested at 40 mg mL^-1^ of total protein concentrated from crude cell extracts. Readings were captured using a nanoliter osmometer and performed in triplicate. Letters represent statistically significant groups following one-way ANOVA with post-hoc Tukey multiple test correction (*p* < 0.01).

**Figure S6.**
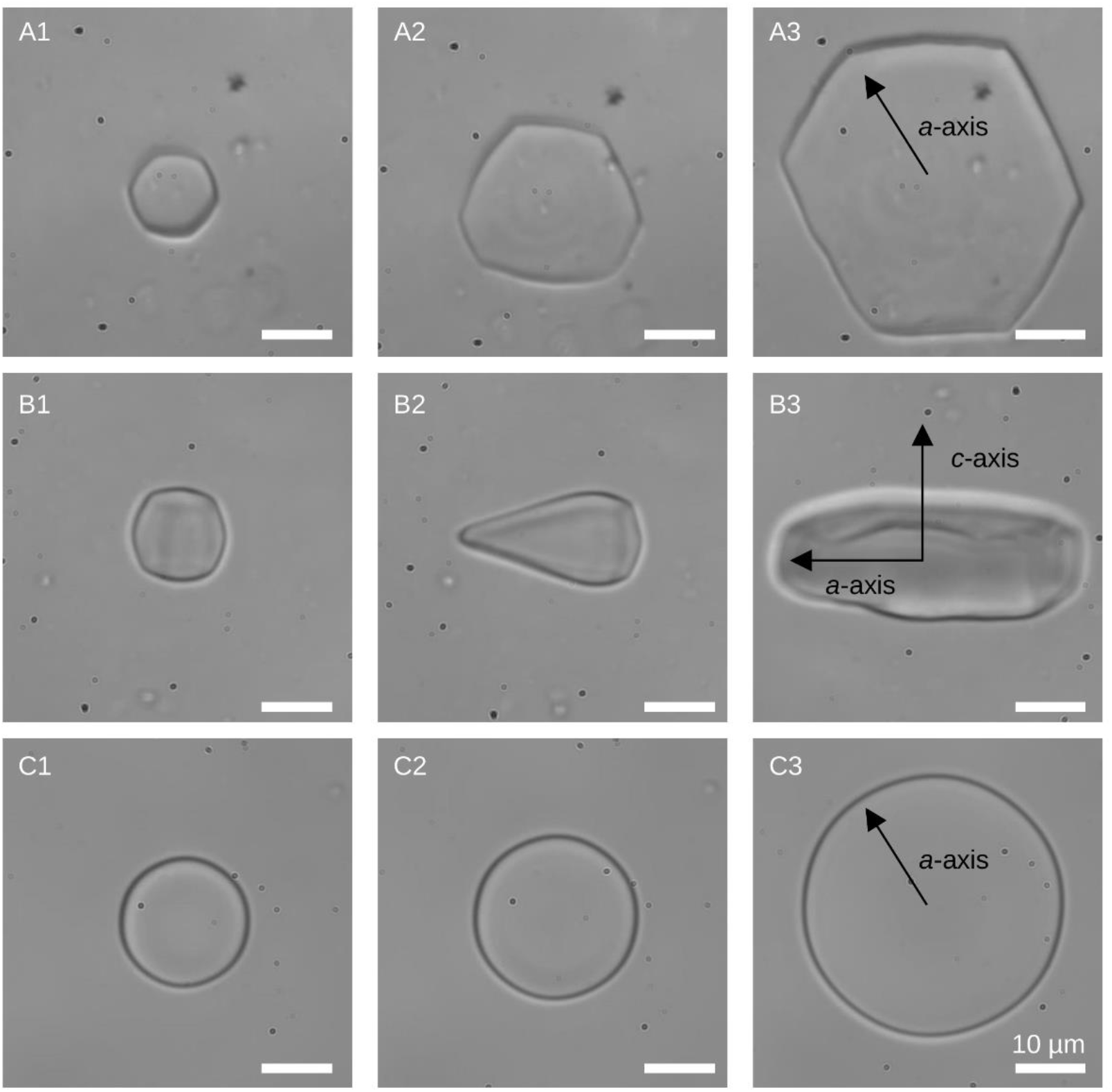
Ice crystal morphology and burst patterns in the presence of antifreeze proteins present in crude lysates of wildtype cold-acclimated *Brachypodium distachyon* Bd21 leaf tissue viewed down on the *a*-axis **(A1-A3)** and viewed horizontally towards the *a-*axis **(B1-B3)**. *Brachypodium* AFPs have an affinity for the prism and basal planes. Buffer solution showing unrestricted ice crystal growth in the absence of AFPs with characteristic disk-shaped morphology visible, viewed down on the *a*-axis **(C1-C3)**. Micrographs were captured at 50x zoom on a nanoliter osmometer and performed in triplicate. Scale bar is 10 µm.

**Figure S7.**
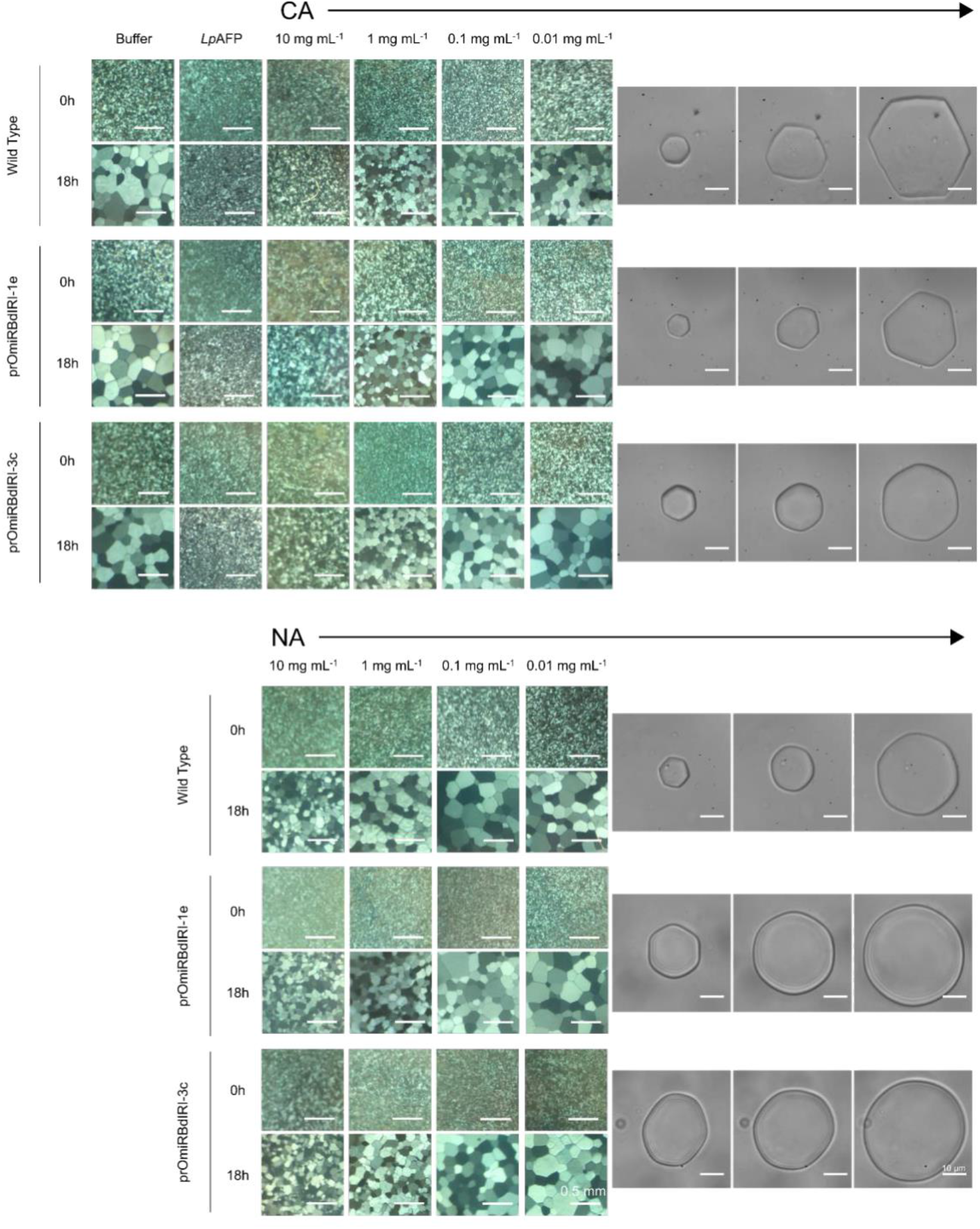
Ice recrystallization inhibition “splat” assays of cell protein extracts using leaf tissue from non-acclimated (NA) and cold-acclimated (CA) *Brachypodium distachyon* Bd21 wild type and temporal AFP knockdown lines prOmiRBdIRI-1e and prOmiRBdIRI-3c. Samples were annealed at - 6 °C for 18 h at various concentrations. Buffer and *Lp*AFP controls are shown. Scale bar for splat assays represents 0.5 mm. Ice crystal morphologies and burst patterns are shown alongside and were tested at 40 mg mL^-1^ of total protein concentrated from crude cell extracts. Micrographs were captured at 50x zoom on a nanoliter osmometer with scale bars representing 10 μM. All assays were performed in triplicate with similar results and representative images are shown.

**Figure S8.**
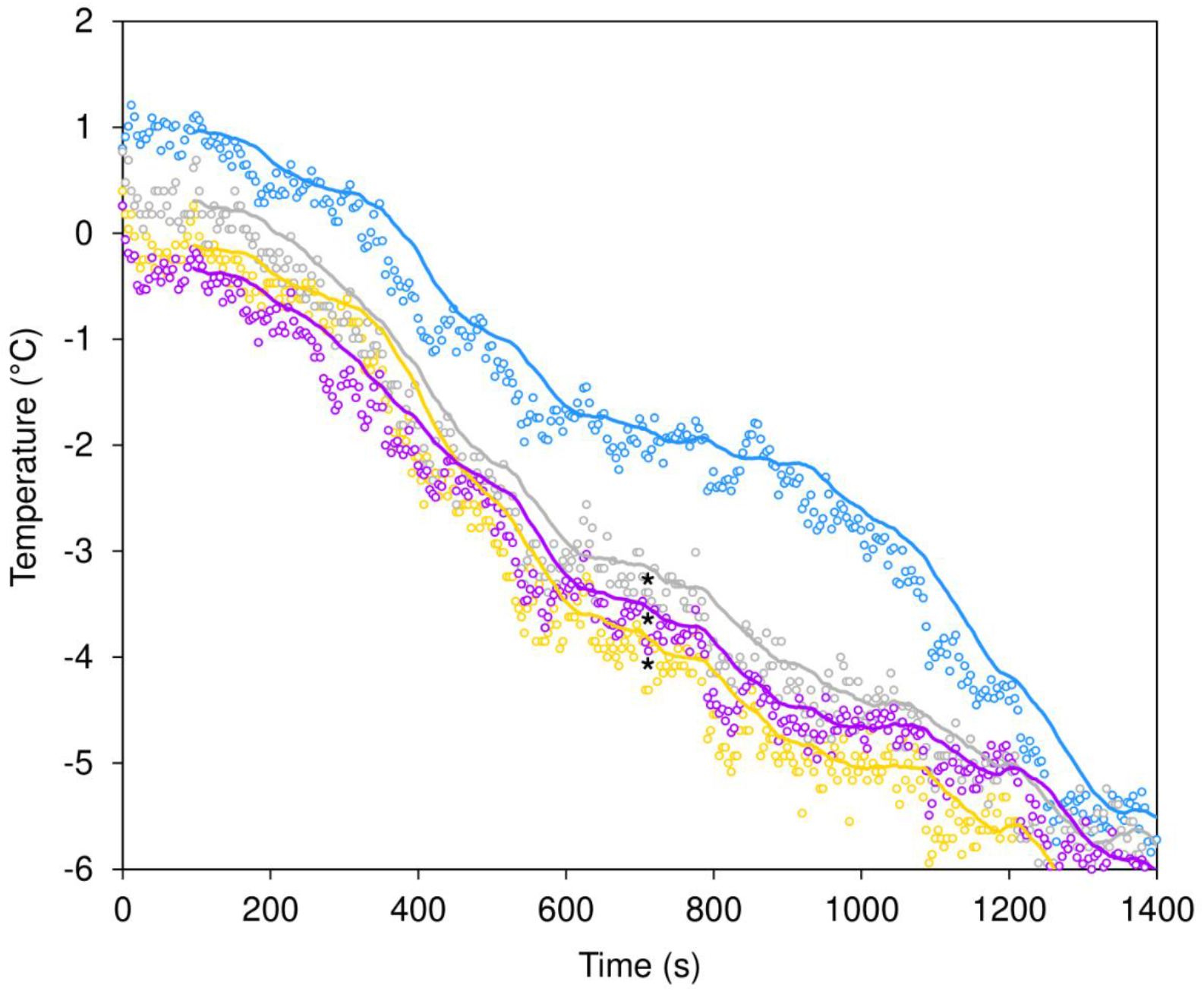
Infrared thermography assay performed on cold-acclimated (CA) wild type (blue), CA prOmiRBdIRI-1e (grey), CA prOmiRBdIRI-3c (purple) knockdown lines, and non-acclimated (NA) wild type (yellow). Data represent the recorded thermography data where each point represents the temperature at a single frame captured at a frame rate of 1 frame every 4 sec. Lines shown represent the moving averages using a period of 25. Points measured were 5 mm from the wounded end of the leaf where the tissue was excised from plants. Leaves were equilibrated at 1 °C for 30 min and frozen to -10 °C at a rate of 0.01 °C sec^-1^. Experiments were performed in triplicate. CA prOmiRBdIRI-1e, CA prOmiRBdIRI-3c knockdown lines, and NA wild type were significantly different from CA wild type indicated by stars (one tailed *t*-test, unpaired).

**Figure S9.**
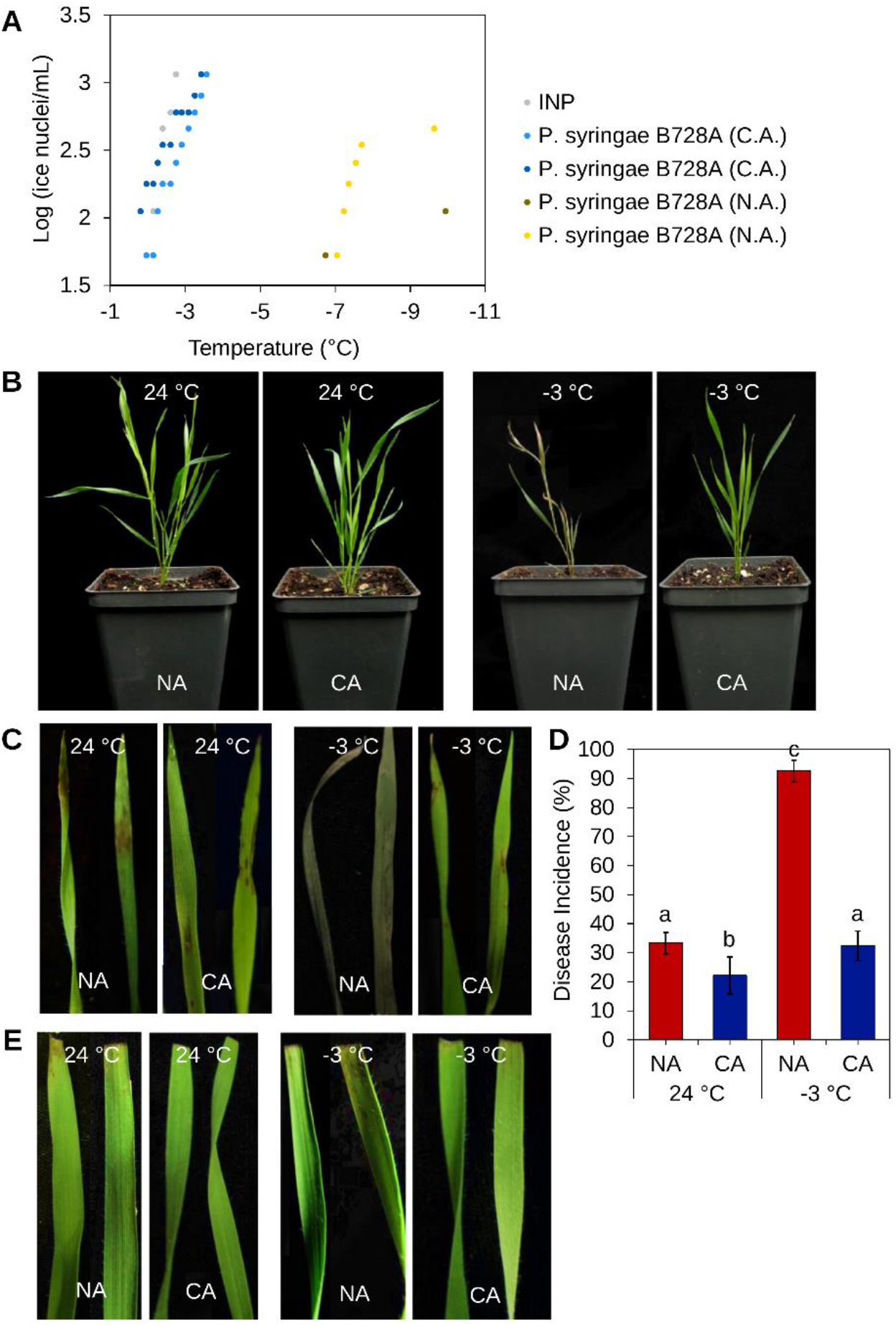
Preliminary experiments for ice nucleation activity and infection protocols of *Brachypodium distachyon* with *Pseudomonas syringae* pv. *syringae* B728A. **(A)** Ice nucleation activity of the *P. syringae* pathovar that had been cold-activated by transfer to 4 °C for 48 h (C.A.) or kept at 24 °C, representing non-cold-activated (N.A.) cultures with nucleation temperatures between -2 °C and -4 °C, were used in assays to determine the conditions and temperature of incubation for leaf tissue in infection assays. *Pseudomonas syringae* ice nucleating protein (INP) preparations (Ward’s Natural Science, Rochester, NY) were used as controls. **(B)** Representative non-acclimated (NA) and cold-acclimated (CA) whole plants sprayed with C.A. *P. syrinage* pathovar cultures and then maintained at 24 °C (two left pots) and -3 °C (two right pots). **(C)** Portions of leaves from NA and CA plants sprayed with C.A. *P. syringae* pathovar cultures at 24 °C standard conditions (the pair of left images) and -3 °C (the pair of right images) showing disease incidence, and in the case of the NA leaves at -3 °C, freeze damage. **(D)** Disease incidence measured as a percentage of leaves sprayed with the C.A. *P. syringae* pathovar showing disease symptoms with leaves from plants that were either NA and CA and then infected as whole plants and kept at the two temperatures shown. **(E)** Leaves taken from NA or CA plants and cut with the wounded ends dipped in cold activated *P. syringae* pathovar culture, with the excised leaves then maintained at 24 °C (left pair of images) and -3 °C (right pair of images).

**Figure S10.**
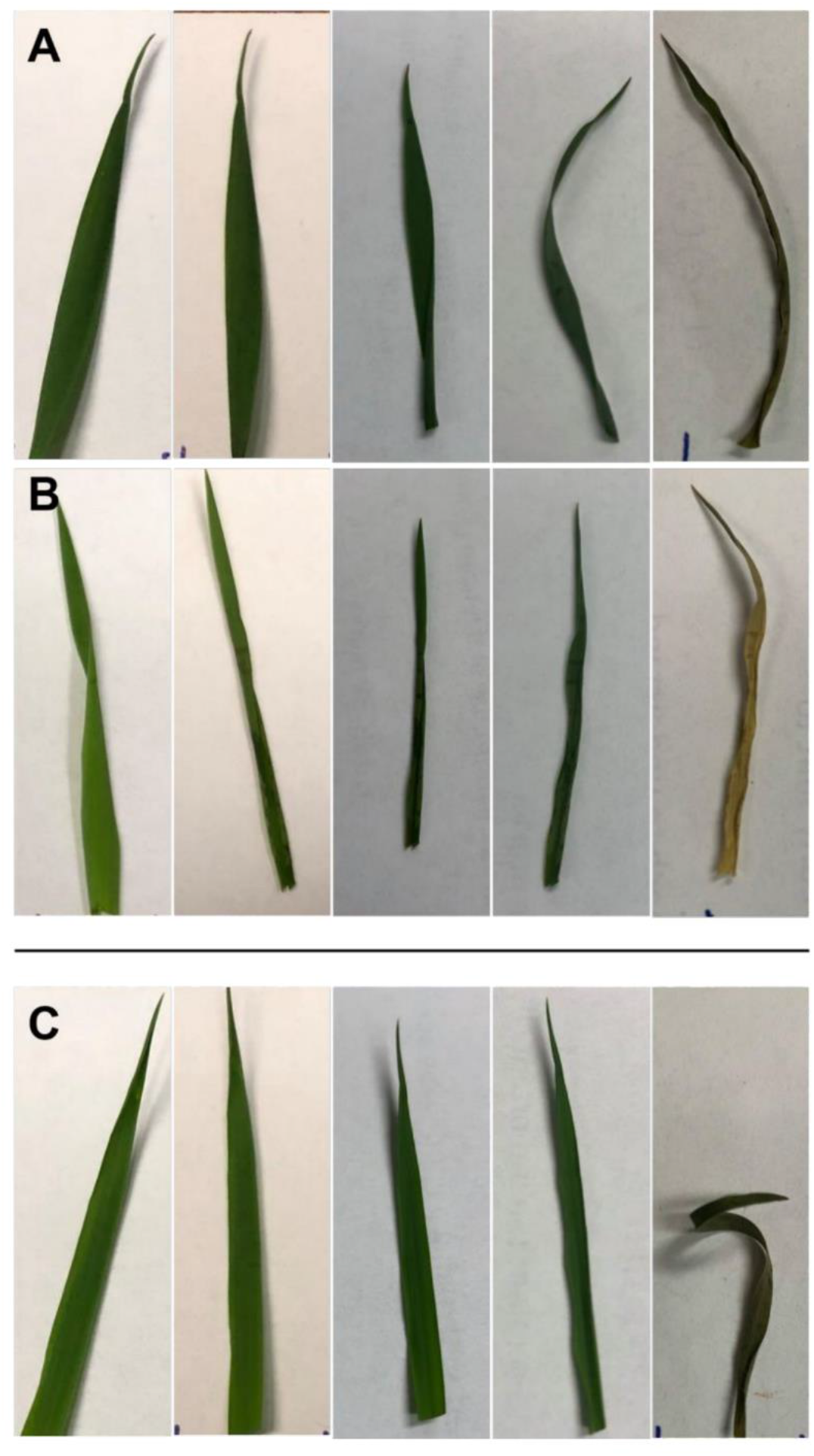
Representative images of excised *Brachypodium distachyon* leaves infected with cold activated *Pseudomonas syringae* pv. *syringae* B728A, with the five images (left to right) representing pre-infection, following infection and incubation for 12 h at -3 °C, and during recovery at 4 °C at 24 h post infection, 48 h post infection, and one week post infection. Samples shown included: **(A)** infected cold-acclimated (CA) Bd21 wild type. **(B)** infected CA temporal cold-induced antifreeze protein (AFP) knockdown line prOmiRBdIRI-1e, and **(C)** uninfected CA Bd21 wild type controls. As indicted in the Methods section, the bacterial strain was cultured at 28 °C to OD_600_ = 0.6-1.0 and placed at 4 °C for two days before resuspending in 10 mM MgCl_2_ and diluted to OD_600_ = 0.2 to an approximate concentration of 1×10^8^ colony forming units mL^-1^ prior to infection by dipping wounded ends of leaves in cultures. Assay was performed in triplicate with similar results and also performed with leaves from non-acclimated plants (not shown).

